# Comparative proteomics of vesicles essential for the egress of *Plasmodium falciparum* gametocytes from red blood cells

**DOI:** 10.1101/2023.02.08.527640

**Authors:** Juliane Sassmannshausen, Sandra Bennink, Ute Distler, Juliane Küchenhoff, Allen M. Minns, Scott E. Lindner, Paul-Christian Burda, Stefan Tenzer, Tim W. Gilberger, Gabriele Pradel

## Abstract

Transmission of malaria parasites to the mosquito is mediated by sexual precursor cells, the gametocytes. Upon entering the mosquito midgut, the gametocytes egress from the enveloping erythrocyte while passing through gametogenesis. Egress follows an inside-out mode during which the membrane of the parasitophorous vacuole ruptures prior to the erythrocyte membrane. Membrane rupture requires the exocytosis of specialized secretory vesicles of the parasites; i.e. the osmiophilic bodies (OBs) involved in rupturing the parasitophorous vacuole membrane, and vesicles (here termed g-exonemes) that harbour the perforin-like protein PPLP2 required for erythrocyte lysis. While several OB proteins are known, like G377 and MDV1/Peg3, the protein composition of the g-exonemes remains unclear. Here, we used high-resolution imaging and BioID methods to study the two types of egress vesicles in *Plasmodium falciparum* gametocytes. We show that OB exocytosis precedes discharge of the g-exonemes and that exocytosis of the g-exonemes, but not of the OBs, is calcium-sensitive. Further, the two types of vesicles exhibit distinct proteomes. In addition to known egress-related proteins, our analyses revealed novel components of OBs and g-exonemes, including proteins involved in vesicle trafficking. Our data provide insight into the immense molecular machinery required for the inside-out egress of *P. falciparum* gametocytes.

## Introduction

The tropical disease malaria is caused by protozoan parasites of the genus *Plasmodium*. The disease is characterized by symptoms like high fever, anemia, nausea, and respiratory distress, mainly caused by the blood-infecting stages of the parasite. Approximately 247 million infections and 619,000 deaths were claimed by malaria in 2021, with two-thirds of the victims being children under five years of age (World Health Organization, 2022). The majority of deaths by malaria are due to infections with *P. falciparum*, the causative agent of malaria tropica.

Malaria is a vector-borne disease with *Plasmodium* parasites being transmitted from human to human by blood-feeding female *Anopheline* mosquitoes. In the human, the parasites follow an obligate intracellular life-style and reside within a parasitophorous vacuole (PV) for most parts of their life-cycle. While vacuolar compartmentalization provides shelter for the parasites to protect themselves from the human immune system, the parasites eventually need to exit the enveloping cell to ensure life-cycle progression (reviewed in Flieger et al., 2018).

To exit their host cells, *Plasmodium* parasites initiate an orchestrated programme of molecular processes, which eventually result in membrane lysis (reviewed in e.g. Wirth & Pradel, 2012; Bennink et al., 2016; Flieger et al., 2018; Sassmannshausen et al., 2020; Bennink & Pradel, 2021; Dvorin & Goldberg, 2022; Tan & Blackman, 2021). Host-cell exit has been particularly studied in the parasite blood stages, i.e., the merozoites and the gametocytes. When egressing from the red blood cell (RBC), the two stages follow an inside-out mode, during which the PV membrane (PVM) ruptures prior to the RBC membrane (RBCM). In both blood stages, the egress cascade begins with the concurrent activation of the plasmodial guanylyl cyclase GCα and the cGMP-dependent protein kinase G following the perception of environmental signals. These events are accompanied by increased intracellular calcium levels, which in consequence activate select calcium-dependent protein kinases. The egress cascade eventually leads to the discharge of specialized secretory vesicles important for the rupture of PVM and RBCM.

In *P. falciparum* merozoites, the egress-related vesicles, termed exonemes, release their content into the PV lumen to initiate RBC lysis. Among others, the exonemes contain the subtilisin-like protease SUB1 and the aspartic protease plasmepsin X (PMX). Once activated by PMX, SUB1 processes a variety of targets that are present in the PV, like the serine-repeat antigen proteins SERA5 and SERA6 as well as the merozoite surface protein MSP1, and these proteins are later required for destabilizing the RBC cytoskeleton prior to RBCM destruction or for mediating the invasion of a new RBC by the freshly released merozoites (Yeoh et al., 2007; Arastu-Kapur et al., 2008; Silmon de Monerri et al., 2011; Ruecker et al., 2012; Stallmach et al., 2015; Collins et al., 2017; Nasamu et al., 2017; Thomas et al., 2018).

Vesicle-mediated exocytosis also plays a crucial role during the egress of gametocytes from RBCs. RBC lysis is mandatory for the gametocytes to finalize gametogenesis, once they are activated in the mosquito midgut by environmental factors. Specialized vesicles of gametocytes, the osmiophilic bodies (OBs), cluster underneath the gametocyte plasma membrane (GPM) within a minute following activation and discharge their content into the PV lumen (Sologub et al., 2011). They were first described in female gametocytes (Sinden, 1982), however, later also male OBs have been reported for *P. berghei*, which exhibit a different protein repertoire and a smaller morphology than the female OBs (Olivieri et al., 2015). A variety of proteins have previously been described that are located in the OBs and released into the PV, the first of which was the female-specific G377 with a to date unknown role in gametogenesis (Alano et al., 1995; Severini et al., 1999; Koning-Ward et al., 2008; Olivieri et al., 2015; Suárez-Cortés et al., 2016). Two other OB-specific proteins, MDV1 (male development gene 1; also termed protein of early gametocytes 3, Peg3) and GEST (gamete egress and sporozoite traversal) are involved in PVM rupture during gametogenesis (Furuya et al., 2005; Ponzi et al., 2009; Silvestrini et al., 2005; Talman et al., 2011; Olivieri et al., 2015; Suárez-Cortés et al., 2016). Moreover, the *P. berghei* gamete egress protein GEP and the putative pantothenate transporter PAT localize to OBs and mutants lacking GEP or PAT are unable to egress from the RBC (Andreadaki et al., 2020; Kehrer et al., 2016b). In addition, several proteases are present in the OBs, like the subtilisin-like protease SUB2 and the dipeptidyl aminopeptidase DPAP2 (Suárez-Cortés et al., 2016). Further, SUB1 and the aspartyl protease MiGS (microgamete surface protein) have been described as MOB-specific proteases, and deficiencies of SUB1 or MiGS result in impaired male gametogenesis (Suárez-Cortés et al., 2016; Tachibana et al., 2018; Pace et al., 2019).

Following secretion of the OB content into the PV lumen, the PVM ruptures at multiple sites before it breaks down into multi-layered vesicles (Sologub et al., 2011). Subsequently, a second vesicle-resident protein is discharged from the activated gametocyte, the *Plasmodium* perforin-like protein PLP2 (Sologub et al., 2011; Deligianni et al., 2013; Wirth et al., 2014). Once secreted, PPLP2 perforates the RBCM, resulting in membrane destabilization and the release of the RBC cytoplasm, while gametocytes deficient of PPLP2 remain trapped in the RBC upon activation (Wirth et al., 2014). A time-span of roughly six minutes was monitored between exocytosis of the OBs and the PPLP2-positive vesicles, indicating that they are two different types of egress vesicles, which act one after the other. Following secretion of PPLP2, it requires another four minutes until the RBC lyses and fully formed gametes are released into the midgut lumen (Sologub et al., 2011; Wirth et al., 2014; Andreadaki et al., 2018).

The data obtained to date point to at least two types of vesicles that are involved in the egress of activated gametocytes from the RBC, i.e. the OBs and the PPLP2-positive vesicles (in the following termed exonemes of gametocytes, g-exonemes). We here aimed to study the dynamics of the OBs and g-exonemes during gametogenesis, to unveil their characteristic components and to identify novel secreted mediators of RBC egress.

## Results

### OBs and g-exonemes differ in their subcellular localization as well as time-point and calcium dependency of discharge

To probe into similarities and differences of OBs and g-exonemes during gametocyte egress we first used immunohistochemical methods to define their subcellular localization and order of discharge. For vesicle visualization, the OB markers G377 and MDV1 and the g-exoneme- resident PPLP2 were chosen. All three proteins display a signal peptide; in addition, PPLP2 possesses a MACPF domain essential for the pore-forming ability (Fig. 1A).

**Figure 1:**
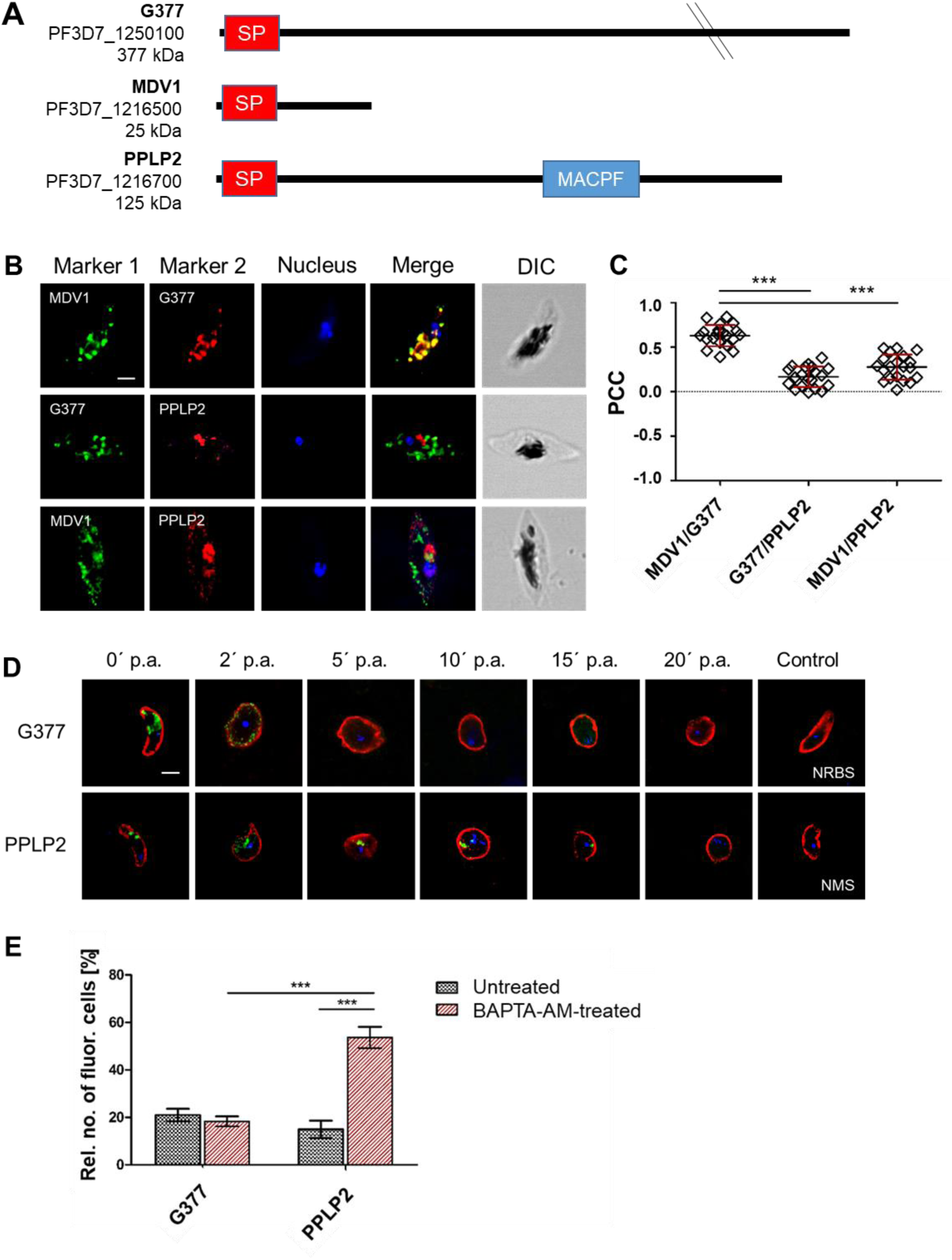
Colocalization and dynamics of egress-related vesicles during gametocyte activation. **(A)** Schematic of the proteins MDV1, G377, and PPLP2. SP, signal peptide; MACPF, Membrane Attack Complex/Perforin domain. **(B)** Vesicular localization of MDV1, G377 and PPLP2 in gametocytes. WT NF54 gametocytes were immunolabeled with rabbit anti-G377, rat anti-MDV1 and mouse anti-PPLP2 antisera (red and green) to investigate co- localization of the respective proteins. Parasite nuclei were highlighted by Hoechst 33342 nuclear stain (blue). Bar; 2 µm. DIC, differential interference contrast. Corresponding negative controls are provided in Fig. S2. **(C)** Quantification of protein colocalization. The Pearsońs correlation coefficient (PCC) was calculated using Fiji ImageJ2. Immunolabeling of G377 or MDV1 was defined as region of interest (n = 20). The error bars indicate mean ± SD. **(D)** Dynamics of G377- and PPLP2-positive vesicles during gametocyte activation. WT NF54 gametocytes were collected at 0-20 min post-activation (p.a.) and immunolabeled, using rabbit anti-G377 or mouse anti-PPLP2 antisera (green). Gametocytes were counterstained with rabbit or mouse anti-P230 antisera (red). Parasite nuclei were highlighted by Hoechst 33342 nuclear stain (blue). Bar; 2 µm. NRBS, neutral rabbit serum; NMS, neutral mouse serum. **(E)** Calcium- dependency of vesicle discharge following gametocyte activation. WT NF54 gametocytes were treated with 25 µM BAPTA-AM prior to activation and immunolabeled with as described in (D). Untreated gametocytes served as control. A total of 50 activated (rounded) gametocytes per setting were evaluated for the presence of the G377 or PPLP2 signal at 20 min p.a. (n = 3). Corresponding IFA images are provided in Fig. S3. The error bars indicate mean ± SD. ***p≤0.001 (One-Way ANOVA with Post-Hoc Bonferroni Multiple Comparison test; C, E). Results (B, D) are representative of three independent experiments.

Indirect immunofluorescence assays (IFAs) were employed to determine the subcellular localization of G377, MDV1, and PPLP2, using specific antisera from rat, rabbit and mouse. All of the proteins were detected in the cytoplasm of wildtype (WT) NF54 gametocytes, localizing to vesicular structures (Fig. 1B; Fig. S1A-C). Both MDV1 and G377 were found in vesicles that are particularly located at the periphery of the gametocytes; PPLP2 was further detected in vesicles accumulating in the central parts of the cell. In accord with previous reports (Severini et al., 1999; Silvestrini et al., 2005; Deligianni et al., 2013; Wirth et al., 2014), G377 and MDV1 are specific for gametocytes and expressed during gametocyte maturation, whereas PPLP2 is present in both the asexual and sexual blood stages (Fig. S1A-C).

Co-labeling IFAs confirmed a colocalization of the two OB markers G377 and MDV1, while the PPLP2 signal did neither overlap with the one of G377 nor of MDV1 (Fig. 1B). No labeling was observed when the gametocytes were immunolabeled with serum of non-immunized mice, rats or rabbits (Fig. S2). Colocalization of the signals were statistically quantified for each double-labeling experiment by calculating the Pearson’s correlation coefficient (PCC). The signal patterns of G377 and MDV1 exhibited a PCC of 0.63, while the PCCs of PPLP2 with G377 or MDV1 had values of 0.17 and 0.28 (Fig. 1C), indicating that the PPLP2-resident g- exonemes are different from the G377- and MDV1-positive OBs.

To investigate the time-points of discharge for the two vesicle types. WT NF54 gametocytes were activated, and samples were taken at several time points between 0 and 20 min post-activation. Subsequently, the two vesicle types were highlighted by immunolabeling of G377 and PPLP2, using the respective antisera (Fig. 1D). While the signal for the OBs disappeared within 2 min post-activation, the PPLP2-resident g-exonemes could be detected for another 10 min, during which the g-exonemes relocated from the center to the periphery of the gametocytes. Around 15 min post-activation, the PPLP2 signal disappeared.

In addition, we determined the role of intracellular calcium as a potential mediator of vesicle discharge. WT NF54 gametocytes were activated in the absence or presence of 25 µM of the selective calcium chelator BAPTA-AM and samples were taken between 0 and 20 min post-activation. The samples were then subjected to IFA, using anti-PPLP2 and anti-G377 antisera to highlight the two vesicle types (Fig. S3). In gametocytes not treated with BAPTA-AM, we again observed that the G377 signal of the OBs disappeared within less than 5 min post-activation, while the PPLP2 signal indicative of g-exonemes relocated to the gametocyte periphery and then disappeared between 10 and 20 min post-activation. When the gametocytes were treated with BAPTA-AM prior to activation, the PPLP2-positive g-exonemes were still detectable at 20 min post-activation, indicating that they were unable to discharge, whereas for G377, no difference in the labeling pattern compared to the untreated control was observed. For quantification, the presence or absence of the PPLP2 and G377 signals at 20 min post-activation was determined in 50 activated rounded gametocytes per experimental setting (Fig. 1E). While the proportion of G377-positive cells was equally low in the presence or absence of BAPTA-AM (18% and 21%, respectively), a significant difference was observed in the proportion of PPLP2-positive cells, which varied between 15% in untreated and 54% in BAPTA-AM-treated gametocytes (Fig. 1E).

Our combined data demonstrate that OBs and g-exonemes are different types of vesicles, which discharge at different time points post-activation with the exocytosis of the OBs preceding discharge of the g-exonemes. Further, we showed that exocytosis of the g-exonemes, but not of the OBs, is sensitive to the calcium chelator BAPTA-AM, indicating its dependency on intracellular calcium.

### OBs and g-exonemes exhibit distinct proteomes with multiple constituents

In order to identify constituents of the OBs and g-exonemes, transgenic parasite lines were generated that express the proteins G377, MDV1 or PPLP2 fused with the promiscuous *E. coli* biotin ligase BirA, which allows the identification of proximal proteins by biotinylation (BioID).

For BioID analyses of putative PPLP2 interactors, a parasite line was generated that episomally expresses PPLP2 fused with BirA and green fluorescent protein (GFP), using the vector pARL- PPLP2-*pffnpa*-GFP-BirA, which expresses the fusion protein under the control of the *pffnpa* promoter (Fig. S4A; (Musabyimana et al., 2022). Furthermore, transgenic lines that endogenously express G377 or MDV1 fused with the enhanced TurboID version of BirA in addition to GFP were generated via homologous recombination using vectors pSLI-G377- TurboID-GFP and pSLI-MDV1-TurboID-GFP, respectively (Fig. S4B; Branon et al., 2018). The episomal presence of vector pARL-PPLP2-*pffnpa*-GFP-BirA in the transgenic parasites and the successful integration of the vectors pSLI-G377-TurboID-GFP and pSLI-MDV1- TurboID-GFP into the respective *g377* or *mdv1* gene loci was demonstrated by diagnostic PCR (Fig. S4C, D).

Western blot analyses of gametocyte lysates prepared from the transgenic lines, using mouse anti-GFP antibody, demonstrated the expression of the respective fusion proteins (Fig. S5A). The proteins MDV1-TurboID-GFP and G377-TurboID-GFP migrated at the expected molecular weights of 91 and 439 kDa, respectively. For PPLP2, a band of approximately 120 kDa was detected in addition to the expected band of 187 kDa, indicating processing of the protein. No protein bands were detected in lysates of either the asexual blood stages of the transgenic lines nor of WT NF54 mixed asexual blood stages and gametocytes (Fig. S5A). Similarly, immunolabeling with anti-GFP antibodies demonstrated the expression of PPLP2- GFP-BirA, MDV1-TurboID-GFP, and G377-TurboID-GFP proteins in gametocytes of the respective transgenic lines and confirmed the presence of the tagged fusion proteins in vesicular structures (Fig. 2A). In WT NF54 parasites, no GFP-labeling was detected.

**Figure 2:**
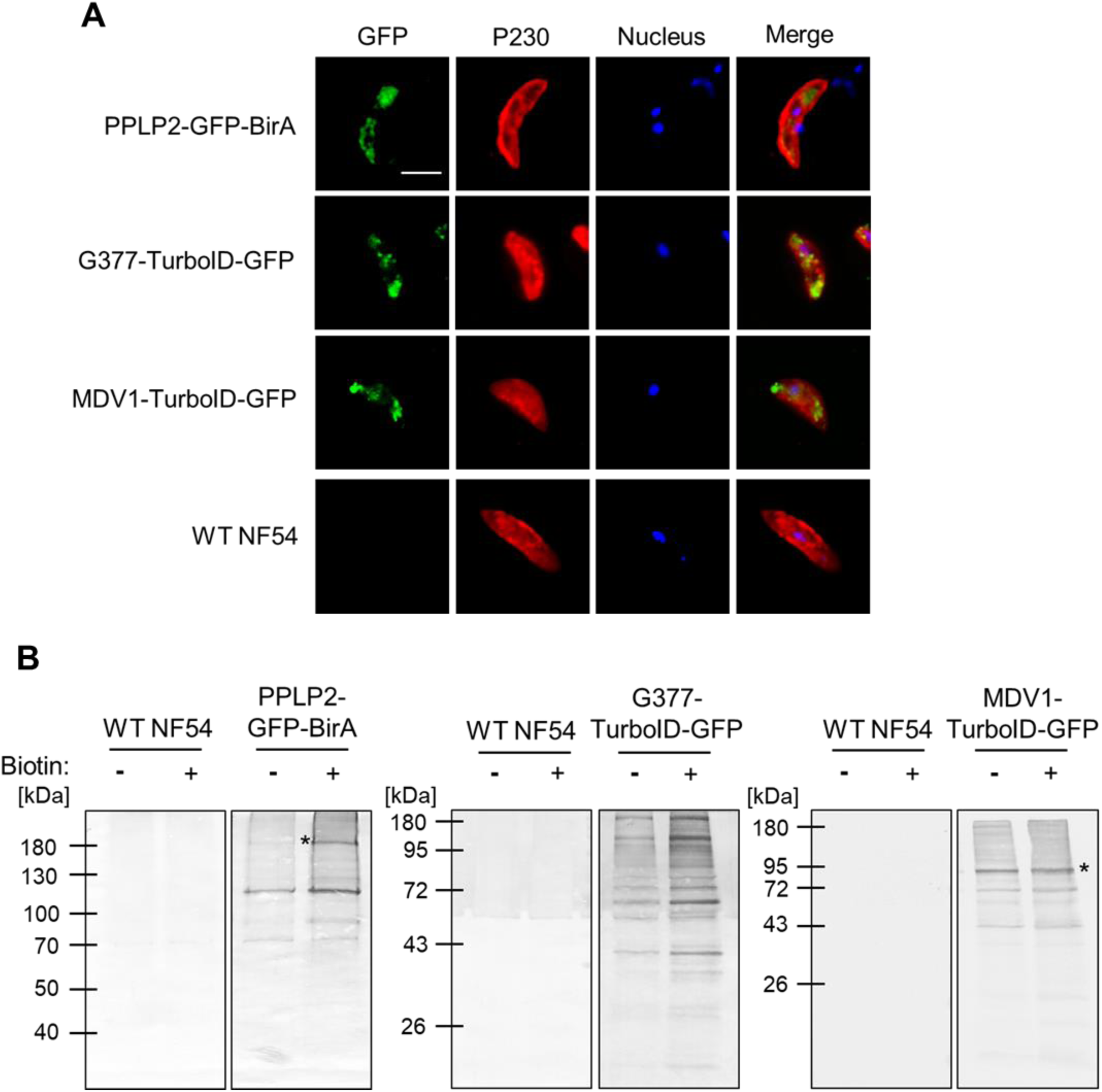
Verification of the parasite lines to be used in BioID. **(A)** Localization of the bait proteins in the BioID parasite lines. Gametocytes of the PPLP2-GFP-BirA line and the G377- TurboID-GFP and MDV1-TurboID-GFP lines were immunolabeled with mouse anti-GFP antibody to highlight PPLP2, G377 and MDV1, fused to GFP and biotin ligase (green). Gametocytes were counterstained with anti-P230 antisera (red); parasite nuclei were highlighted by Hoechst 33342 nuclear stain (blue). WT NF54 gametocytes served as a control. Bar; 5 µm. **(B)** Protein biotinylation in the BioID parasite lines. Gametocytes of the PPLP2- GFP-BirA line and the G377-TurboID-GFP and MDV1-TurboID-GFP lines were treated with biotin (+) for 20 h (PPLP2-GFP-BirA) and 15 min (G377-, MDV-TurboID-GFP). Untreated parasite lines (-) and biotin-treated and untreated NF54 WT gametocytes served as controls. Gametocyte lysates were subjected to Western blot analysis and biotinylated proteins were detected using streptavidin-conjugated AP. Asteriks (*) highlight the bait proteins. Cloning strategy and transfection verification are provided in Fig. S4. Results (A, B) are representative of three independent experiments.

Subsequent Western blotting was employed to highlight biotinylated proteins in the transgenic lines. For this, gametocyte cultures of each line were treated with 50 µM biotin for 15 min (TurboID) or 20 h (BioID). Immunoblotting of the respective gametocyte lysates using streptavidin conjugated to alkaline phosphatase detected multiple protein bands indicative of biotinylated proteins, including protein bands of 91 kDa and 187 kDa, likely representing the biotinylated fusion proteins MDV1-TurboID-GFP and PPLP2-GFP-BirA, respectively, while the fusion protein G377-TurboID-GFP could not be identified uniquely due to the high molecular weight (Fig. 2B). In gametocytes of the transgenic lines that were not treated with biotin, minor protein bands were detected, indicating endogenous biotin has been present in the gametocytes, which triggered the activity of the biotin ligase. No biotin-positive protein bands were detected in WT NF54 samples (Fig. 2B). IFA analyses of biotin-treated gametocytes of the transgenic lines, using fluorophore-conjugated streptavidin, confirmed the presence of biotinylated proteins, which localized in vesicular structures, while no biotinylated proteins were detected in biotin-treated WT NF54 gametocytes (Fig. S5B).

BioID analyses were subsequently employed to analyze the proteomes of the OBs and g- exonemes of *P. falciparum*. For this, gametocytes of the respective transgenic lines were treated with biotin as described above, and equal amounts of gametocytes per sample were harvested. Three independent samples were collected from each of the three lines (MDV1- TurboID-GFP, G377-TurboID-GFP and PPLP2-GFP-BirA); two additional independent samples from the PPLP2-GFP-BirA line were included. Mass spectrometric analysis was performed on streptavidin-purified protein samples with three technical replicas for each sample. This resulted in the identification of 636 (MDV1-TurboID-GFP), 189 (G377-TurboID-GFP), and 298 (PPLP2-GFP-BirA) significantly enriched proteins, respectively (Fig. 3A; Table S1). For each transgenic parasite line, the respective bait protein, i.e. G377, MDV1, and PPLP2 was detected among these proteins (marked in Table S1). In a first analysis step proteins without a putative signal peptide and/or transmembrane domains (with no C-terminal ER retention signal) were excluded, reducing the potential interactors to 169 proteins (MDV1- TurboID-GFP), 50 proteins (G377-TurboID-GFP), and 64 proteins (PPLP2-GFP-BirA) (Fig. 3A; Table S2). Subsequently, proteins with defined known functions not related to OBs or g- exonemes (e.g. nucleoporins, chaperons) were removed from the list, eventually resulting in the following numbers of putative proteins of egress vesicles: 132 proteins (MDV1-TurboID-GFP), 38 proteins (G377-TurboID-GFP), and 44 proteins (PPLP2-GFP-BirA) (Fig. 3A; Table S2).

**Figure 3:**
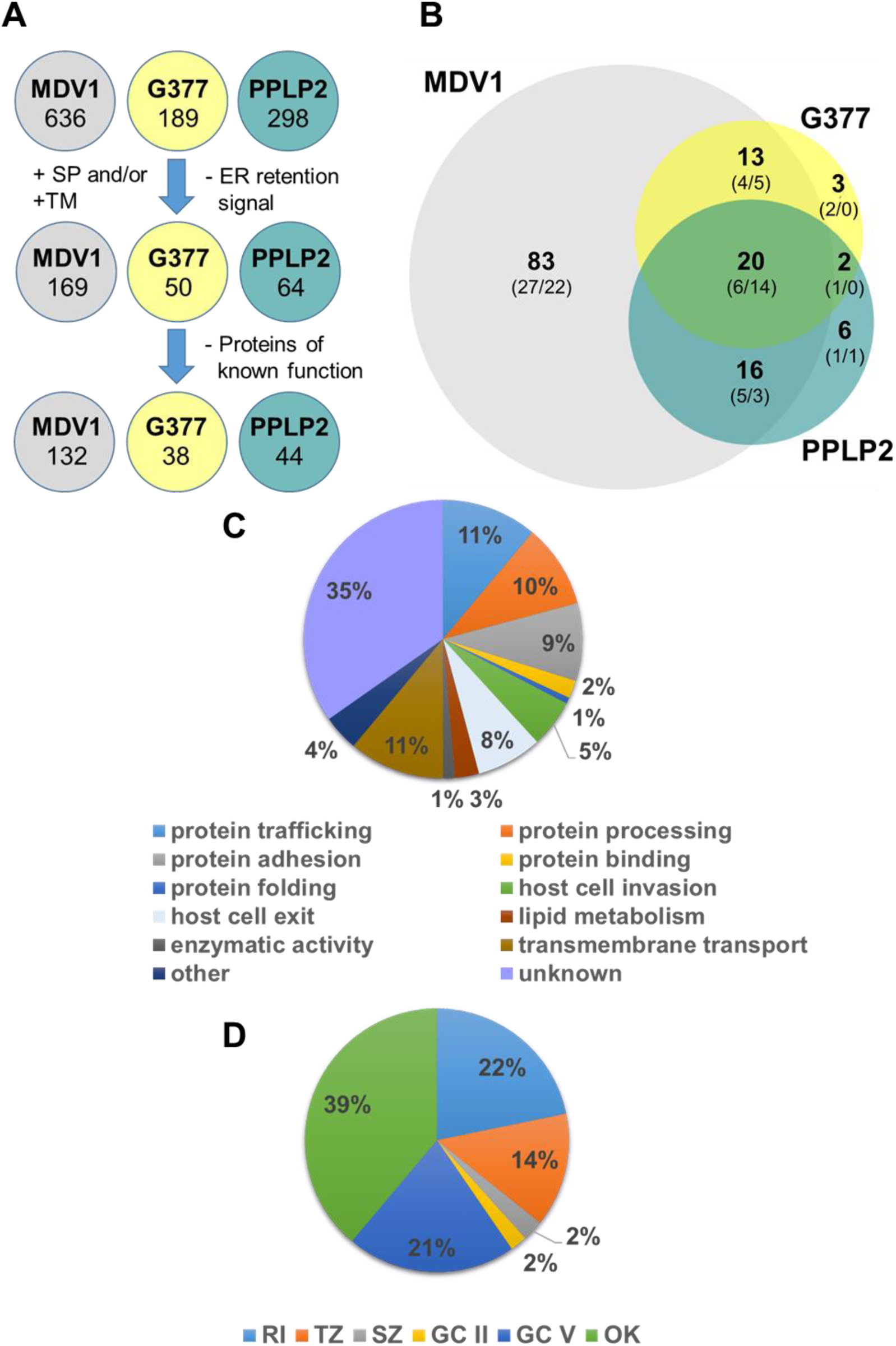
*In silico* analysis of the egress vesicle proteomes. **(A)** Schematic of candidate selection. Putative interactors of MDV1, G377, and PPLP2 following BioID of lines MDV1- TurboID-GFP, G377-TurboID-GFP and PPLP2-GFP-BirA were subjected to domain and functional analysis, resulting in the identification of a total of 143 egress vesicle proteins. Signal peptides (SP) were predicted using SignalP 4.1 & 5.0, transmembrane domains (TM) were predicted using DeepTMHMM, and endoplasmic reticulum (ER) retention signals were predicted using DeepLoc 2.0. Functional prediction was performed via PlasmoDB. **(B)** Venn diagram depicting the egress vesicle proteins grouped by bait protein. The final numbers of interactors for each parasite line are depicted in bold, the predicted sex specificity is indicated (male/female; Lasonder et al., 2016; see PlasmoDB database). **(C)** Pie chart depicting the egress vesicle proteins (percentage of total numbers) according to predicted molecular function. **(D)** Pie chart depicting the egress vesicle proteins (percentage of total numbers) grouped by stages of peak expression (López-Barragán et al., 2011; see PlasmoDB database). RI, ring stage; TZ, trophozoite; SZ, schizont; GC II, gametocyte stage II; GC V, gametocyte stage V; OK, ookinete. Detailed information on the interactors before and after the application of selection criteria is provided in Tables S1 and S2. The corresponding GO term and sex specificity analyses are provided in Fig. S6.

The comparison of the filtered lists of proteins identified 13 proteins as putative interactors of both OB proteins, G377 and MDV1. It also revealed 16 proteins as putative interactors of MDV1 and PPLP2, while only 2 proteins were shared between G377 and PPLP2 (Fig. 3B; Table S2). In total, 20 proteins were shared between three bait proteins. This group of proteins includes among others five members of the LCCL protein family, as well as P230 and P230p, and P47, hence adhesion proteins known to locate in the PV of gametocytes, where they form protein complexes that are linked to the GPM (Simon et al., 2009, 2016; reviewed in Kuehn et al., 2010).

The putative interactors were then grouped by predicted function (Fig. 3C). The majority of proteins belonged to the categories of protein trafficking, processing and adhesion as well as transmembrane transport (∼10 % in each category). A total of 8% of proteins was previously assigned to host cell exit, e.g. EPF1 (exported protein family 1), GEP (gamete egress protein), GEXP02 (gametocyte exported protein 2), PMX (plasmepsin X), SUB2 (subtilisin-like protease 2), MiGS (microgamete surface protein), and GEST (gamete egress and sporozoite traversal protein) (Talman et al., 2011; Kehrer et al., 2016a; Suárez-Cortés et al., 2016; Nasamu et al., 2017; Tachibana et al., 2018; Andreadaki et al., 2020; Grasso et al., 2022). Further, 35% of proteins are of unknown function. An additional gene ontology (GO) term analysis revealed main molecular functions in peptidase activities and pyrophosphate hydrolysis and well as transmembrane transport (Fig. S6A) and cellular localizations in host cellular components and vesicles (Fig. S6B).

A comparative transcriptional analyses of these putative interactors (according to table “Transcriptomes of 7 sexual and asexual life stages”; López-Barragán et al., 2011; see PlasmoDB database; Aurrecoechea et al., 2009) showed that the majority of proteins exhibited peaks in stage V gametocytes and in ookinetes or in ring stages and trophozoites (Fig. 3D), suggesting that they represent two groups of proteins, either present during gametocyte development, e.g. P230, P48/45, or with roles in the mosquito midgut phase, e.g. members of the PSOP family. When the sex specificity of the interactors was evaluated (according to table “Gametocyte Transcriptomes”; Lasonder et al., 2016; see PlasmoDB database; Aurrecoechea et al., 2009), roughly one-third of proteins could be assigned to either male or female gametocytes, while one third of proteins did not exhibit sex-specific transcript expression (Fig. S6C).

The interactors were further subjected to STRING-based analyses to investigate the protein-protein interaction networks (see string-db.org; text mining included). The STRING analysis revealed three main clusters (Fig. 4; Table S3). The first cluster involved proteins previously shown to form multi-adhesion domain protein complexes like the LCCL domain proteins, P48/45, and P230, plus the paralogs P47 and P230p (Pradel et al., 2004; Scholz et al., 2008; Simon et al., 2009; 2016; reviewed in Kuehn et al., 2010; Pradel, 2007). Furthermore, the cluster includes three members of the PSOP family, PSOP1, PSOP12 and PSOP13 (Ecker et al., 2008; Sala et al., 2015; Tachibana et al., 2021) and proteins linked to the PVM, i.e. P16, Pfg17-744 and Pfg14-748 (Bruce et al., 1994; Eksi et al., 2005). In addition, the three vesicle markers G377, MDV1 and PPLP2, are found in this cluster. Noteworthy, the majority of these proteins are interactors of all three bait proteins.

**Figure 4:**
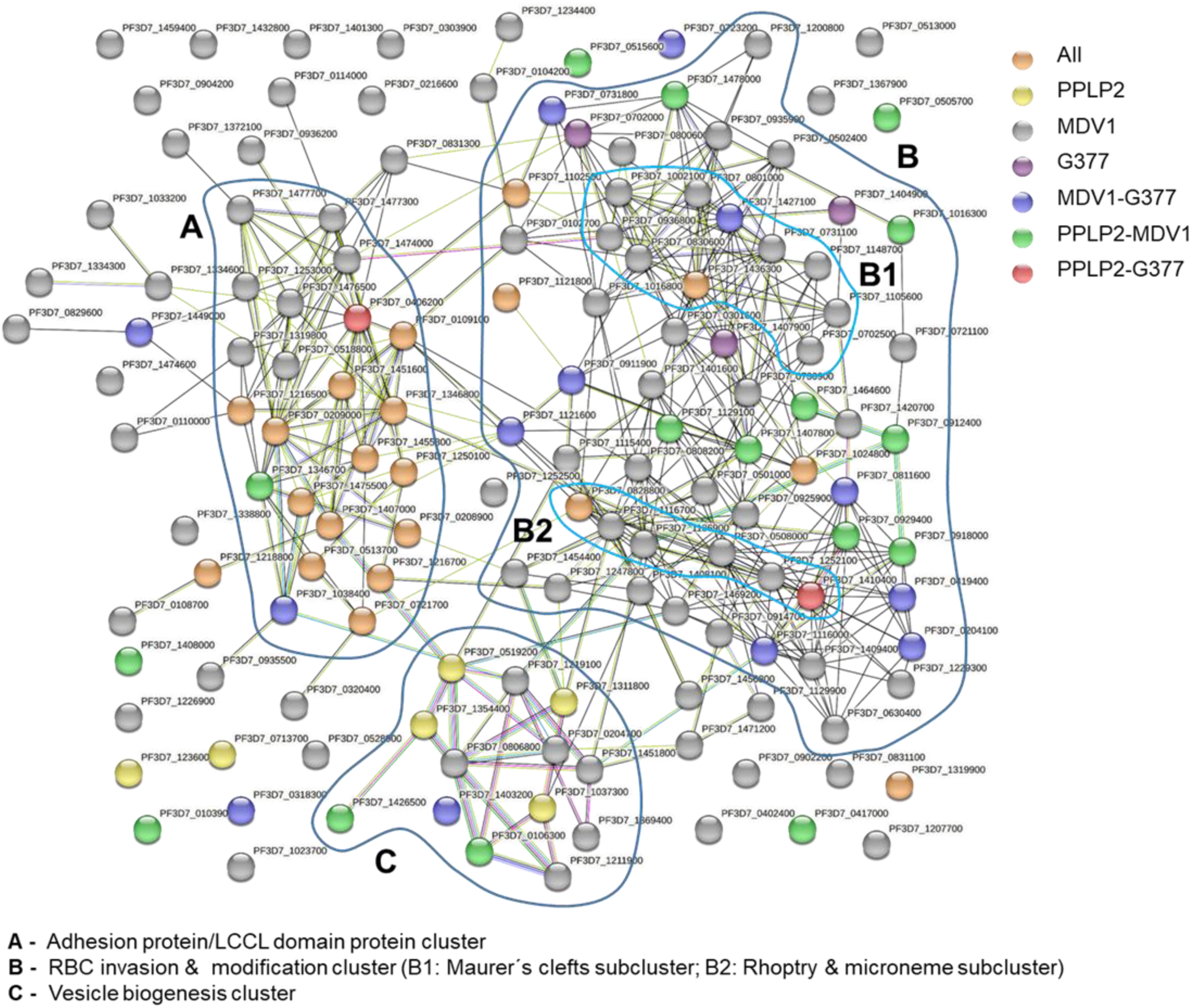
Network analysis of the egress vesicle proteins. The 143 egress vesicle proteins were evaluated for potential interactions using the STRING database. Based on the interaction among the query proteins, different functional clusters were identified. **(A)** Adhesion protein/LCCL domain protein cluster with 23 proteins; **(B)** the RBC invasion and modification cluster with 59 proteins, including **(B1)** the Maureŕs clefts subcluster and **(B2)** the rhoptry/microneme subcluster; **(C)** the vesicle biogenesis cluster of 13 proteins. Detailed information on individual proteins clustering in each subnetwork is provided in Table S3.

A second cluster comprises proteins linked to vesicle biogenesis, particularly transmembrane transporters including the previously described ABCG2 transporter of female gametocytes (Tran et al., 2014). Further, the vesicle trafficking-related protein clathrin (heavy chain) and sortilin are found in this cluster (Fig. 4; Table S3). Four of the proteins of this cluster belong to the group of PPLP2 interactors.

The third cluster resembles a megacluster that comprises various proteins particularly linked to RBC invasion and modification. Within the megacluster, two subclusters can be distinguished, one of which includes proteins linked to the Maureŕs clefts, while the other one comprises proteins linked to rhoptries and micronemes. Additional proteins are associated with these subclusters, several of which are proteases, e.g. SUB2, falstatin, the dipeptidyl aminopeptidases DPAP1 and DPAP2, the metalloprotease M16, and the plasmepsins PMI, PMIII, PMX. Furthermore, known components of the merozoite surface like GAMA, MaTrA, MSP8 or P38 as well as various exported proteins like GEXP08, GEXP21, EXP1 and EXP3 are found within the megacluster (Fig. 4, Table S3). Since the majority of the megacluster proteins were previously linked to rings and trophozoites, they may have functions in both the asexual and sexual blood stages.

The fact that several known proteins where found in both the OB and g-exoneme interactomes suggests that these may have met during endomembrane trafficking. In this context, we validated the vesicular localization of CCp2, a component of the LCCL domain adhesion protein complex known to be synthesized continuously and present at the GPM (reviewed in Kuehn et al., 2010; Pradel, 2007). CCp2 as well as other components of the adhesion protein complex have been identified as interactors of all three bait proteins (see above). IFA analyses confirmed the presence of CCp2 in vesicles and in association with the GPM as well as the plasma membrane of gametes, where it neither co-localizes with G377 nor PPLP2 (Fig. S7).

In conclusion, we identified various proteins as potential constituents of egress vesicles. Among the candidates, previously described components of OBs and g-exonemes as well as novel proteins were found. Novel candidates particularly included peptidases, transmembrane transporters and proteins involved in vesicle trafficking.

### Selected putative constituents localize to vesicular structures in gametocytes

Out of the 143 putative constituents of egress vesicles, five proteins previously not studied in gametocytes were chosen from the different groups of interactors for further validation, i.e. the Sel1-repeat containing protein (PF3D7_0204100), the secreted ookinete protein PSOP1 (PF3D7_0721700) and an unknown conserved protein (PF3D7_0811600), which interact with G377 and MDV1, the PPLP2-interacting v-SNARE protein Vti1 (PF3D7_1236000) of g- exonemes and an unknown conserved protein (PF3D7_1319900), which was identified as an interactor of G377, MDV1 and PPLP2. Transgenic lines were generated, using vector pSLI- HA-*glmS* (Musabyimana et al., 2022), which leads to the expression of the protein of interest fused to a hemagglutinin A (HA)-tag in addition to the *glmS* ribozyme in the 3’ untranslated region (Fig. S8A). Successful vector integration into the targeted locus was shown by diagnostic PCR (Fig. S8B).

Western blotting of lysates derived from all of the five HA-tagged parasite lines confirmed its expression in the *P. falciparum* blood stages. Four of the HA-tagged proteins were detected in lysates of both asexual blood stages and gametocytes, running at the expected molecular weights, i.e. Sel1 (271 kDa), PSOP1 (53 kDa), PF3D7_0811600 (144 kDa), Vti1 (49 kDa; Fig. 5A). The conserved protein PF3D7_1319900 was restricted to gametocyte lysates and in addition to the expected 179 kDa protein, a second band running at approximately 120 kDa was identified, indicating protein processing. No protein bands were detected in lysates of WT parasites, which served as a control (Fig. 5A).

**Figure 5:**
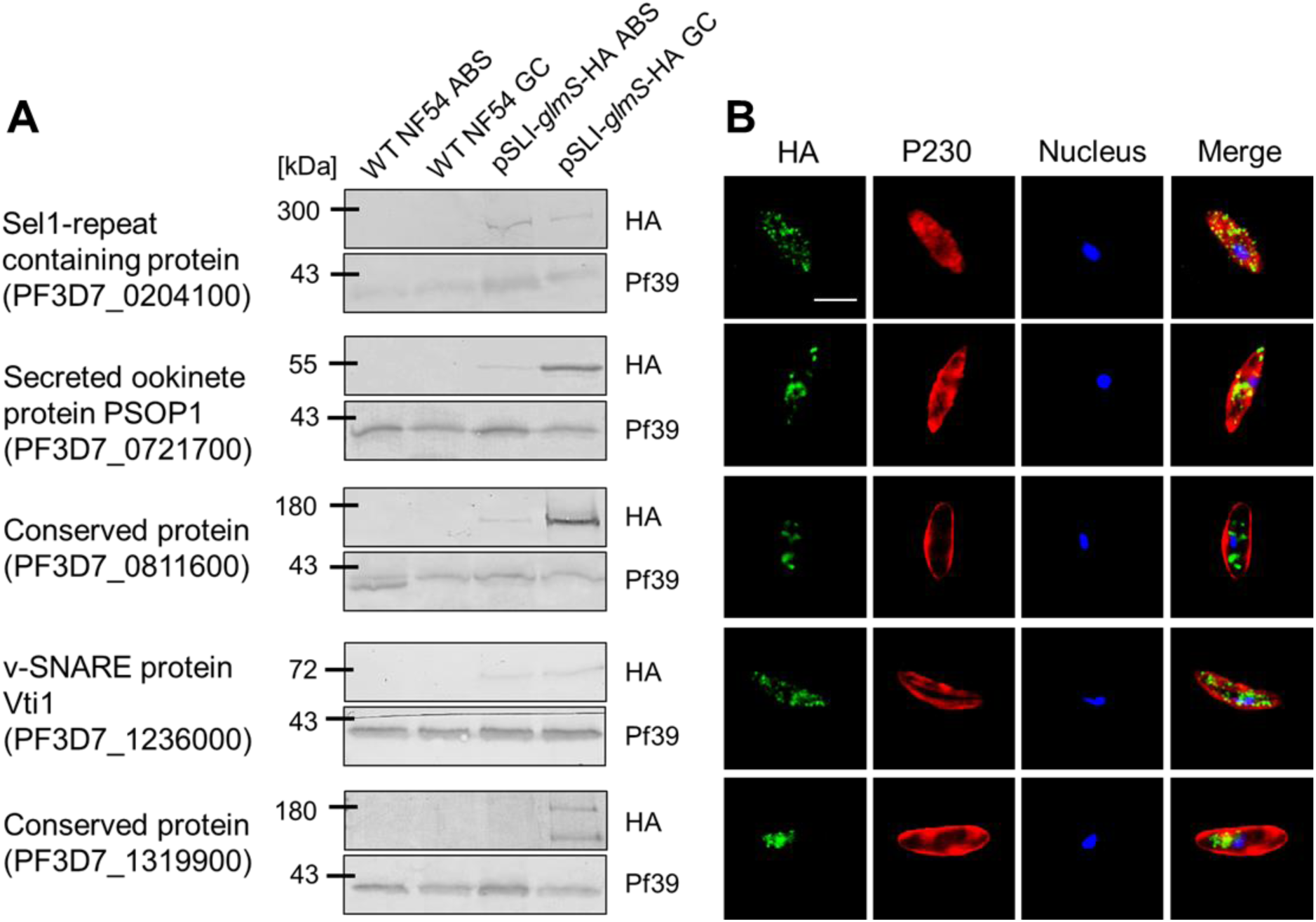
Expression and localization of egress vesicle proteins in blood stage parasites. Five pSLI-HA-*glmS*-based parasite lines expressing select egress vesicle proteins tagged with HA were generated for expression analysis. **(A)** Blood stage expression of egress vesicle proteins. Lysates of asexual blood stages (ABS) and gametocytes (GC) of the respective lines were immunoblotted with rat anti-HA antibody to detect Sel1 (271 kDa), PSOP1 (53 kDa), the conserved protein PF3D7_0811600 (144 kDa), Vti1 (49 kDa) and the conserved protein PF3D7_1319900 (179 kDa). ABS and GC lysate of WT NF54 served as negative controls, while immunoblotting with rabbit antibody against the ER protein Pf39 (39 kDa) served as loading control. **(B)** Vesicular localization of egress vesicle proteins in gametocytes. Gametocytes of the pSLI-HA-*glmS*-based parasite lines were immunolabeled with rat anti-HA antibody (green) to detect the respective HA-tagged egress vesicle protein. Gametocytes were counterstained with anti-P230 antisera (red); parasite nuclei were highlighted by Hoechst 33342 nuclear stain (blue). Bar; 5 µm. Cloning strategy and transfection verification are provided in Fig. S8. Results (A, B) are representative of three independent experiments.

The subcellular localization of the HA-tagged proteins in gametocytes was confirmed by IFAs. All of the proteins showed a punctuate localization within the gametocyte cytoplasm, clearly defining vesicular structures (Fig. 5B). While for the majority of proteins, the vesicular pattern was distributed over the cell body and also found in the cell periphery, the conserved protein PF3D7_1319900 was observed mainly in the center of the gametocytes, indicating that the protein might be restricted to the ER-Golgi complex.

For the HA-tagged PSOP1 and the unknown conserved protein PF3D7_0811600, we evaluated the vesicular localization in more detail via super-resolution microscopy. Co-labeling experiments, using anti-HA antibodies, revealed overlapping signals of both PSOP1 and PF3D7_0811600 with G377, while the two proteins did not co-label with PPLP2 (Fig. 6A). No labeling was observed, when gametocytes of the respective PSOP1-HA-*glmS* and PF3D7_0811600-HA-*glmS* lines were immunolabeled with serum of non-immunized mice, rats or rabbits (Fig. S9). Colocalization was quantified in IFAs and PCC values were evaluated for each double-labeling experiment. The signal patterns of PSOP1 and PF3D7_0811600 overlapped with G377 with PCC values of 0.85 and 0.6 respectively, pointing to a localization of the two proteins in the OBs. In contrast, the signal overlap of the two proteins with PPLP2 resulted low in PCC values of 0.33 and 0.29, respectively (Fig. 6B).

**Figure 6:**
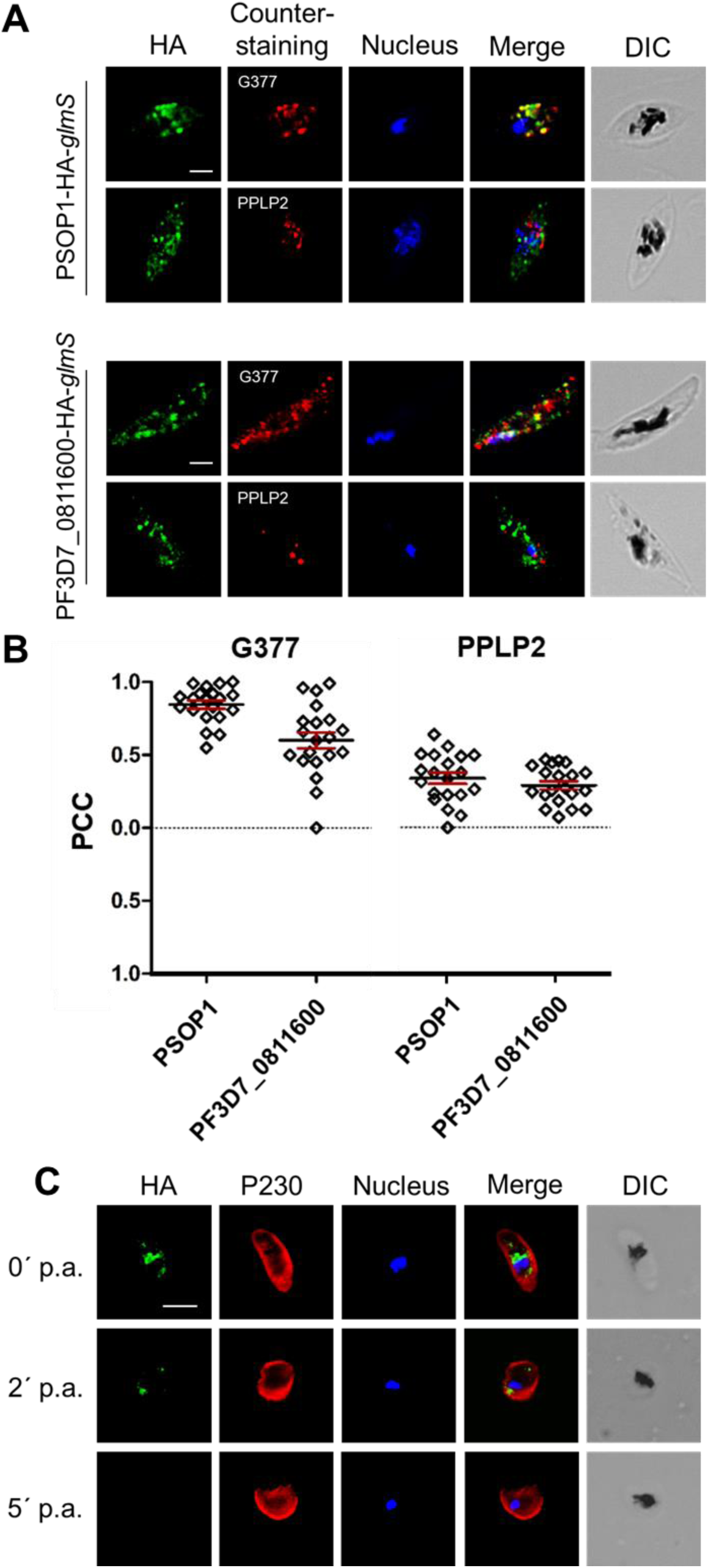
Localization and discharge of OB components. **(A)** Localization of PSOP1 and PF3D7_0811600 in OBs. Gametocytes of the PSOP1-HA-*glmS* line and PF3D7_0811600-HA- *glmS* line were immunolabeled with rat anti-HA antibody (green). OBs and g-exonemes were highlighted using rabbit anti-G377 or mouse anti-PPLP2 antisera (red); parasite nuclei were highlighted by Hoechst 33342 nuclear stain (blue). Samples were analyzed via Airyscan super- resolution microscopy. Bar; 2 µm. DIC, differential interference contrast. **(B)** Quantification of protein colocalization. The Pearsońs correlation coefficient (PCC) was calculated using Fiji ImageJ2. Immunolabeling of HA-tagged egress proteins was defined as region of interest (n = 20). The error bars indicate mean ± SD. **(C)** PSOP1 discharge following gametocyte activation. Gametocytes of line PSOP1-HA-*glmS* were collected at 0-5 min post-activation (p.a.) and immunolabeled with rat anti-HA antibody to highlight PSOP1 (green). Gametocytes were counterstained by rabbit anti-P230 antisera (red); parasite nuclei were highlighted by Hoechst 33342 nuclear stain (blue). Bar; 5 µm. DIC, differential interference contrast. Results (A, C) are representative of three independent experiments.

Eventually, the fate of PSOP1 during gametocyte activation was determined. Activated gametocytes between 0 and 20 min were immunolabeled with anti-HA antibody to detect PSOP1. The gametocytes were either counterstained by P230 labeling or the OBs were highlighted by G377 immunolabeling (Fig. 6C; Fig. S10A, B). The IFAs revealed that both PSOP1 and G377 were discharged simultaneously within the first two minutes following activation.

In conclusion, we confirmed the vesicular expression of selected candidates of OBs and g- exonemes and in addition demonstrated that PSOP1 is a marker of OBs in *P. falciparum* gametocytes, which is released into the PV lumen immediately following gametocyte activation.

## Discussion

The exit of activated gametocytes from the RBC is mediated by molecules that are discharged by specialized secretory vesicles following the perception of the egress signal. Activation triggers the exocytosis of OBs needed to rupture the PVM as well as of recently identified PPLP2-positive g-exonemes that are important for the perforation of the RBCM.

Two previous proteomics analyses were performed, using the rodent malaria model *P. berghei*, to unveil on the egressome of gametocytes (Kehrer et al., 2016a; Grasso et al., 2020). In both approaches, the entity of proteins released during gametogenesis was collected and determined. In addition, a MDV1-specific interactome was established. The two studies described a variety of secreted proteases, like serine-rich antigens SERA1, SERA2, SERA3, the subtilisin-like proteases SUB1, SUB2, MiGS, and PMX as egressome components. Additional proteins included GEST, MDV1, PPLP2, G377, MSP1 and MSP9, and MTRAP as well as PSOP1, PSOP12 and PSOP17.

The aim of the present study was to evaluate the types and proteomes of gametocyte-specific egress vesicles in the human malaria species *P. falciparum*. We firstly demonstrated that OBs and g-exonemes are two different types of vesicles and described the consecutive exocytosis of the two vesicle types following gametocyte activation, with the latter exocytosis being dependent on intracellular calcium. These observations are in accord with our previous findings reporting that OBs discharge at 1 min post-activation, while PPLP2-resident vesicles discharge approximately 6 min later and that PPLP2 release, but not OB exocytosis, can be impaired by calcium chelators (Sologub et al., 2011; Wirth et al., 2014).

We then analyzed interactors of the OB-specific proteins G377 and MDV1 and of the g- exoneme component PPLP2 in order to characterize the gametocyte egressome. On the one hand, we found several previously described mediators of egress, including ones identified in the above mentioned proteomics analysis of egressing *P. berghei* gametocytes. These egress proteins comprise, in addition to the bait proteins, among others Pf11-1, GEST, GEP, MiGS, GEXP02, SUB2, EPF1, and PMX (Scherf et al., 1992; Talman et al., 2011; Mbengue et al., 2013; Kehrer et al., 2016a; Olivieri et al., 2015; Suárez-Cortés et al., 2016; Nasamu et al., 2017; Pino et al., 2017; Tachibana et al., 2018; Pace et al., 2019; Andreadaki et al., 2020; Grasso et al., 2020; Warncke et al., 2020; reviewed in Pradel, 2007; Kuehn & Pradel, 2010; Bennink & Pradel, 2021). Noteworthy, the latter two were so far assigned to egressing merozoites, but appear to play further roles in RBC exit by gametocytes. On the other hand, we identified novel constituents or proteins previously assigned to other functions and lifecycle stages, e.g. merozoite proteins like RhopH2, MaTrA, MSP8, and P38, all of which are known to be involved in RBC invasion (Curtidor et al., 2004; Ntumngia et al., 2004; Black et al., 2005; Sanders et al., 2005; Vincensini et al., 2008; Hinds et al., 2009; reviewed in Cowman et al., 2017; Molina-Franky et al., 2022), or the PSOP members PSOP1, PSOP12, PSOP13, PSOP17 and GAMA (the ortholog of *P. berghei* PSOP9) originally described in the ookinete micronemes and assigned to mosquito midgut penetration (Ecker et al., 2008; Tachibana et al., 2021). The fact that the here identified egressome includes proteins important for merozoite invasion, as well as ookinete proteins assigned to midgut extravasation, suggests that such proteins may have multiple functions related to host cell membrane modification in different lifecycle stages.

We also identified various proteases, e.g. SUB2, falstatin, DPAP1, DPAP2, M1AAP, M16, as well as PMI, PMIII, PMX, the majority of which were so far linked to hemoglobin processing (reviewed in e.g. Rosenthal, 2002; Goldberg, 2005; Trenholme et al., 2010; Siqueira-Neto et al., 2018; Arisue et al., 2020; Nasamu et al., 2020). Importantly, a high number of further interactors was previously associated to the Maureŕs clefts, like the gametocyte exported protein family members GEXP02, GEXP17, GEXP12, GEXP21, and GEXP22, further EPF1, PTP5, ETRAMP10.2, MSRP5, and PV1 as well as various PHIST proteins and unnamed exported proteins. We hypothesize that these proteins function in remodeling the erythrocyte cytoskeleton as a preceding step of RBCM rupture.

Noteworthy, the majority of the identified proteins were interactors of MDV1, which could be assigned, in equal parts, to male and female gametocytes or were considered to be sex-unspecific. MDV1 was originally described to function during male gametogenesis (Furuya et al., 2005); however, other reports neither confirmed its male-specificity in natural infections (Ponzi et al., 2009; Tadesse et al., 2019) or assigned it to female gametocytes (Lal et al., 2009). This discrepancy in the sex assignment of MDV1 would explain, why no link between its interactors and any potential male-specific expression could be detected.

The identification of PPLP2-specific interactors proved difficult, even after repetition of the BioID analysis using another independent duplicate. The majority of PPLP2 interactors can be assigned to vesicle biogenesis, particularly transmembrane transporters. To be highlighted is the interaction of PPLP2 with the v-SNARE protein Vti1, which is one of three identified *P. falciparum* orthologs of eukaryotic Vti1 (Ayong et al., 2007; Aurrecoechea et al., 2009).

Indeed, *in silico* and transcript and protein expression analyses confirmed that many factors of the vesicle trafficking and fusion machinery, including SNAREs and associated proteins, are conserved in *P. falciparum* and expressed in gametocytes (reviewed in Bennink & Pradel, 2021). The role of these molecules in the dynamics of egress vesicles, however, is yet unknown and needs to be investigated in more detail.

Several of the identified candidates were interactors shared by PPLP2 with G377 and/or MDV1. The majority of these proteins belong to the LCCL domain protein-based multi- adhesion domain protein complex, including CCp1, CCp2, CCp3, CCp5 and FNPA as well as the associated proteins P48/45, and P230, plus the paralogs P47 and P230p. It has previously been shown that the components of this multi-adhesion domain protein complex are expressed throughout gametocyte maturation and localize in the PV, where the complex is anchored via P48/45 in the GPM (Pradel et al., 2004; Scholz et al., 2008; Simon et al., 2009, 2016; reviewed in Pradel, 2007; Kuehn et al., 2010). IFAs confirmed the presence of CCp2 in vesicles and the PV and showed that CCp2 does neither co-label with G377 nor PPLP2. These results suggest that the components of the multi-adhesion domain protein complex have met the biotin ligase-tagged bait proteins in the secretory pathway, hence during endomembrane trafficking. This would also explain why PPLP2 was previously identified as an interactor of MDV1 (Kehrer et al., 2016a).

In a last step of our study, we selected five known and yet uncharacterized components of the egress vesicle interactomes, including the novel v-SNARE Vti1, and verified their vesicular localization. In addition, we confirmed an OB-specific localization of PSOP1 and the unknown gene product PF3D7_0811600. The verification of PSOP1 as an OB-resident protein strengthens our hypothesis that PSOP proteins have additional functions distinct from ookinete biology. In this context, a recent study identified naturally acquired antibodies against PSOP1 in blood samples of various malaria cohorts, verifying the presence of the protein in gametocytes (Muthui et al., 2021). Similarly, the presence of PSOP12, another egress molecule identified in this study that is a member of the 6-cys family, was demonstrated to be expressed in gametocytes in addition to ookinetes (Wass et al., 2012; Annoura et al., 2014; Sala et al., 2015). The expression of such PSOP members from gametocyte maturation until ookinete development makes them promising candidates of transmission blocking vaccines (Sala et al., 2015).

In conclusion, we demonstrated the presence of two different types of egress vesicles with important functions during RBC exit by activated gametocytes. We further verified the existence of various OB components in *P. falciparum*, which were previously identified in *P. berghei*. In addition, we identified novel components of the two egress vesicle types, including Maureŕs cleft-associated proteins, proteases, transmembrane transporters, and vesicle trafficking proteins, but also various yet unknown proteins. Follow-up studies will need to verify the role of these candidates in gametogenesis in order to shed more light on the molecular machinery of the inside-out egress of gametocytes.

## Experimental Procedure

### Gene Identifiers

Aldolase (PF3D7_1444800), CCp2 (PF3D7_1455800); Conserved protein #1 (PF3D7_0811600); Conserved protein #2 (PF3D7_1319900); G377 (PF3D7_1250100); MDV1 (PF3D7_1216500); Pf39 (PF3D7_1108600); Pf92 (PF3D7_1364100); P230 (PF3D7_0209000); PPLP2 (PF3D7_1216700), PSOP1 (PF3D7_0721700); Sel1 (PF3D7_0204100), Vti1 (PF3D7_1236000).

### Bioinformatics

Colocalization of G377, MDV1 and PPLP2 in IFAs was evaluated via Pearsońs correlation coefficient (PCC) using Fiji ImageJ2 (Schindelin et al., 2012). Prediction of the transmembrane domains and intra-/extracellular regions of the putative interaction partners was performed using DeepTMHMM (Hallgren et al., 2022); further signal peptide prediction was performed using the SignalP versions 4.1 with sensitive D-cutoff and 5.0 (Petersen et al., 2011; Almagro Armenteros et al., 2019). Subcellular localization regarding potential ER localization was predicted using DeepLoc 2.0 (Külzer et al., 2009; Thumuluri et al., 2022). Predictions of gene expression and protein function were made using the database PlasmoDB (http://plasmoDB.org; Aurrecoechea et al., 2009); the peak transcript expression of candidate genes was analyzed using table “Transcriptomes of 7 sexual and asexual life stages” (López-Barragán et al., 2011) and sex specificity was predicted using table “Gametocyte Transcriptomes” (Lasonder et al., 2016) of the PlasmoDB database. A sex specificity was considered, when the transcript per million (TPM) ratio was ≥ 3. The gene ontology enrichment (GO) analysis was performed using the PlasmoDB database with a p-value cut-off of 0.05. A network analysis was conducted using the STRING database (version 11.0) (Szklarczyk et al., 2019), using default settings, including text mining options and a confidence of 0.009.

### Antibodies

The following antisera were used: rabbit polyclonal antisera against G377 (Severini et al., 1999; kindly provided by Pietro Alano, Istituto Superiore di Sanità, Rome, Italy), CCp2 (Simon et al., 2009, 2016; Roling et al., 2022), Pf39 (Simon et al., 2009), Pf92 (Musabyimana et al., 2022) and P230 (Simon et al., 2009, 2016); mouse polyclonal antisera against Pf92 (Musabyimana et al., 2022); PPLP2 (Wirth et al., 2014), CCp2 (Pradel et al., 2004), or P230 (Williamson et al., 1995); rat polyclonal antisera against MDV1 (kindly provided by Pietro Alano, Istituto Superiore di Sanità, Rome, Italy), monoclonal mouse anti-GFP antibody (Roche; Basel, CH) and monoclonal rat anti-HA antibody (Roche; Basel, CH).

### Parasite Culture

In this study, *P. falciparum* strain NF54 (WT NF54) was used. The WT NF54 and all generated mutant parasite lines were cultivated *in vitro* in human blood group A+ erythrocytes as previously described (Ifediba & Vanderberg, 1981). Asexual blood stages and gametocytes were maintained in RPMI1640/HEPES medium (Gibco; Thermo Fisher Scientific; Waltham, US) supplemented with 10% (v/v) heat inactivated human A+ serum, 50 μg/ml hypoxanthine (Sigma-Aldrich; Taufkirchen, DE) and 10 μg/ml gentamicin (Gibco; Thermo Fisher Scientific; Waltham, US). For cultivation of the mutant parasite lines, the selection drug WR99210 (Jacobus Pharmaceutical Company; Princeton, US) was added in a final concentration of 2.5 nM or 4.0 nM. All cultures were kept at 37°C in an atmosphere of 5% O_2_ and 5% CO_2_ in N_2_. Gametocytes were enriched via Percoll (Cytiva; Washington DC, US) gradient centrifugation as described previously (Kariuki et al., 1998). Gametogenesis was induced by adding xanthurenic acid in a final concentration of 100 μM dissolved in 1% (v/v) 0.5 M NH_4_OH/ddH_2_O and incubation for 15 min at room temperature (RT). To analyze the role of intracellular calcium during exocytosis, the cultures were treated with 25 µM BAPTA-AM (Thermo Fisher Scientific; Waltham, US) for 15 min at 37°C prior to activation.

### Generation of parasite lines to be used in BioID

Plasmid pSLI-TurboID-GFP (pSL1423) was created to serve as a base plasmid for the fusion of TurboID with the C-terminus of the respective protein-of-interest. The previously described pSLI-TGD plasmid (Addgene #85791, Birnbaum et al., 2017) was modified by inserting a recodonized version of TurboID (“V4”) and a portion of GFPmut2 between the existing NotI and HpaI sites. Vector pSLI-TurboID-GFP (pSL1423) will be made available for request through Addgene.

For the generation of the MDV1- and G377-TurboID-GFP parasite lines, a gene fragment homologous to the 3’-region of the gene was amplified via PCR using the respective primers (for primer sequences, see Table S4). Ligation of insert and vector backbone was mediated by NotI and SpeI restriction sites. A NF54 WT culture with at least 5% ring stages was transfected with 100 µg plasmid DNA in transfection buffer via electroporation (310 V, 950 μF, 12 ms; Bio-Rad gene-pulser Xcell) as described (e.g. Wirth et al., 2014; Ngwa et al., 2017). At 6 h post-transfection, WR99210 was added in a final concentration of 4 nM. A mock control was electroporated using transfection buffer without the addition of plasmid DNA and was cultured in medium with and without WR99210. After approximately 3-5 weeks, WR-resistant parasites appeared in the cultures. To remove NF54 WT parasites from the culture, these were treated with 800 µg/ml neomycin (G418 disulfate salt; Sigma-Aldrich; Taufkirchen, DE) daily for a maximum of 7 days. To verify successful integration into the respective gene locus, genomic DNA (gDNA) was isolated from the transgenic parasite lines, using the NucleoSpin Blood Kit (Macherey-Nagel; Dueren, DE) following the manufacturer’s protocol and used as template in diagnostic PCR. The following primers were used to confirm vector integration: 5’Int MDV1/G377-TurboID, 3’Int MDV/G377-TurboID, pARL-HA-*glmS* FP, pGREP RP (for primer sequences, see Table S4).

The PPLP2-GFP-BirA parasite line was generated by using vector pARL-*pffnpa*-GFP-BirA (Musabyimana et al., 2022). The full-length gene was amplified via PCR using the respective primers (for primer sequences, see Table S4). Ligation of insert and vector backbone was mediated by KpnI and AvrII restriction sites. Transfection of the construct was carried out as described above. Episomal presence of the construct was checked by using the following primers: PPLP2-BirA-KpnI-FP and pGREP RP (for primer sequences, see Table S4).

### Generation of pSLI-HA-*glmS* parasite lines

The HA-*glmS*-tagged parasite lines were generated via single-crossover homologous recombination, using vector pSLI-HA-*glmS* (Musabyimana et al., 2022; kindly provided by Dr. Ron Dzikowski, The Hebrew University of Jerusalem, Israel). A gene fragment homologous to the 3’-region of the gene was amplified via PCR using the respective primers (for primer sequences, see Table S4). The ligation of insert and vector backbone was mediated by NotI and XmaI restriction sites. The transfection and selection procedures were performed as described above, but WR99210 was added in a final concentration of 2.5 nM. The following primers were used to confirm vector integration: 5’-Integration Construct pSLI-*glmS*, 3’-Integration Construct pSLI-*glmS*, pARL-HA-*glmS* FP, pSLI-HA-*glmS* RP (for primer sequences, see Table S4).

### Western Blot Analysis

Asexual blood stage parasites of WT NF54, lines MDV1- and G377-TurboID-GFP, line PPLP2-GFP-BirA and the HA-*glmS*-tagged parasite lines were harvested from mixed cultures. For erythrocyte lysis, parasites were incubated for 10 min in 0.05% (w/v) saponin/PBS. Gametocytes were enriched by Percoll purification as described above. Pelleted parasites were resuspended in lysis buffer (150 mM NaCl, 0.1% (v/v) Triton X-100, 0.5% (w/v) sodium deoxycholate, 0.1% (w/v) SDS, 50 mM Tris-HCl, pH 8.0) supplemented with protease inhibitor cocktail (PIC, Roche; Basel, CH). A 5x SDS loading buffer with 25 mM dithiothreitol was added to the lysates, followed by heat-denaturation for 10 min at 95°C. Lysates were separated via SDS-PAGE and transferred to a nitrocellulose membrane. Non-specific binding sites were blocked by incubation with 5% (w/v) skim milk and 1% (w/v) BSA in Tris-buffered saline (pH 7.5) for 1 h at 4°C. For immunodetection, membranes were incubated overnight at 4°C with monoclonal mouse anti-GFP antibody (Roche; Basel, CH; dilution 1:500) or monoclonal rat anti-HA antibody (Roche; Basel, CH; dilution 1:200) and polyclonal rabbit anti-Pf39 antisera (dilution 1:10,000) in 3% (w/v) skim milk/TBS. The membranes were washed 3x each with 3% (w/v) skim milk/TBS and 3% (w/v) skim milk/0.1 % (v/v) Tween/TBS and then incubated for 1 h at RT with goat anti-mouse, anti-rabbit or anti-rat alkaline phosphatase-conjugated secondary antibodies (dilution 1:10,000, Sigma-Aldrich; Taufkirchen, DE) in 3% (w/v) skim milk/TBS. The membranes were developed in a NBT/BCIP solution (nitroblue tetrazolium chloride/5-bromo-4-chloro-3-indoxyl phosphate; Roche; Basel, CH) for up to 20 min at RT. For the detection of biotinylated proteins, the blocking step was performed overnight at 4°C in 5% (w/v) skim milk/TBS and the membrane was washed 5x with 1x TBS before incubation with streptavidin-conjugated alkaline phosphatase (dilution 1:1,000, Sigma-Aldrich; Taufkirchen, DE) in 5% (w/v) BSA/TBS for 1 h at RT.

### Indirect Immunofluorescence Assay

Asexual blood stage parasites and gametocytes of WT NF54, lines MDV1- and G377-TurboID-GFP, line PPLP2-GFP-BirA, and the HA-*glmS*-tagged parasite lines were coated on diagnostic slides (Epredia; Thermo Fisher Scientific; Waltham, US) and air-dried. Fixation of the cells was performed with 40 μl of 4% (w/v) paraformaldehyde (pH 7.2) for 10 min at RT, followed by membrane permeabilization with 30 μl of 0.1% (v/v) Triton X-100/125 mM glycerol for another 10 min at RT. Non-specific binding sites were blocked by incubation with 3% (w/v) BSA/PBS for 30 min. The primary antibodies were diluted in 3% (w/v) BSA/PBS and were added to the slide for 2 h incubation at 37°C. The slides were washed 3x with PBS and incubated with the secondary antibody for 1 h at 37°C. Following 2x washing with PBS, the incubation with the second primary antibody and the corresponding visualization with the second secondary antibody were carried out as described above. The nuclei were stained with Hoechst 33342 staining solution for 2 min at RT (1:5,000 in 1x PBS). The cells were mounted with anti-fading solution (Citifluor Limited; London, UK), covered with a coverslip and sealed airtight with nail polish. The parasites were visualized by either conventional fluorescence microscopy using a Leica DM5500 B (Leica; Wetzlar, DE) or by confocal microscopy using Zeiss LSM880 Airyscan super-resolution microscope (Carl Zeiss; Jena, DE) fitted with an Airyscan detector and a Plan-Apochromat x63 (NA 1.4) M27 oil objective. During confocal microscopy, the images were acquired sequentially with a pixel resolution of 0.02 µm in four channels as follows: channel 1 = 405 nm laser, channel 2 = 488 nm laser, channel 3 = 561 nm laser, channel 4 = differential interference contrast (DIC). Antibodies were diluted as follows: rat anti-MDV1 (1:1,000), rabbit anti-G377 (1:1,000), mouse anti-PPLP2 (1:50), mouse anti-GFP (1:500), rat anti-HA (1:100), anti-CCp2 (mouse, 1:50; rabbit, 1:500), mouse and rabbit anti-Pf92 (1:500) mouse and rabbit anti-P230 (1:400), and anti-mouse Alexa Fluor 488, anti-rabbit Alexa Fluor 488, anti-rat Alexa Fluor 488, anti-mouse Alexa Fluor 594, anti-rabbit Alexa Fluor 594, anti-mouse Alexa Fluor 555 and anti-rabbit Alexa Fluor 555 (1:1,000; used for LSM880 microscopy; all fluorophores from Invitrogen Molecular Probes; Eugene, US); further Alexa Fluor 594 streptavidin (1:500; Invitrogen Molecular Probes; Eugene, US) was used.

### Statistical analysis

Data are expressed as mean ± SD. Statistical differences were determined using One-Way ANOVA with Post-Hoc Bonferroni Multiple Comparison test. P-values <0.05 were considered statistically significant. Significances were calculated using GraphPad Prism 5 and are defined as follows: *p<0.05; **p<0.01; ***p<0.001.

### Affinity purification of biotinylated proteins

Biotinylation was induced by treatment of the corresponding gametocyte cultures with 50 µM biotin for 15 min (TurboID) or 20 h (BioID) at 37°C. Gametocytes were washed in RPMI incomplete medium, enriched via Percoll gradient centrifugation, and resuspended in 100 µl binding buffer (Tris-buffered saline containing 1% (v/v) Triton X-100 and protease inhibitor). The sample was sonicated on ice (2 x 60 pulses at 30% duty cycle) and another 100 µl of icecold binding buffer was added. After a second session of sonification, cell debris was pelleted by centrifugation (5 min, 16,000x *g*, 4°C). The supernatant was mixed with pre-equilibrated Cytiva Streptavidin Mag Sepharose Magnet-Beads (Cytiva; Washington DC, US) in a low-binding reaction tube. Incubation was performed with slow end-over-end mixing over night at 4°C. The beads were washed 6x with 500 μl washing buffer (3x: RIPA buffer containing 0.03% (w/v) SDS, followed by 3x 25 mM Tris buffer, pH 7.5) and were resuspended in 100 μl elution buffer (1% (w/v) SDS/5 mM biotin in Tris buffer (pH 7.5)), followed by an incubation for 5 min at 95°C. The supernatant was transferred into a new reaction tube and stored at 4°C.

### Proteolytic digestion

Samples were processed by single-pot solid-phase-enhanced sample preparation (SP3) as described before (Hughes et al., 2014; Sielaff et al., 2017). In brief, proteins bound to the streptavidin beads were released by incubating the samples for 5 min at 95° in an SDS-containing buffer (1% (w/v) SDS, 5 mM biotin in water/Tris, pH 8.0). After elution, proteins were reduced and alkylated, using DTT and iodoacetamide (IAA), respectively. Afterwards, 2 µl of carboxylate-modified paramagnetic beads (Sera-Mag Speed Beads, GE Healthcare; Chicago, US; 0.5 μg solids/μl in water as described by Hughes et al., 2014) were added to the samples. After adding acetonitrile to a final concentration of 70% (v/v), samples were allowed to settle at RT for 20 min. Subsequently, beads were washed twice with 70% (v/v) ethanol in water and once with acetonitrile. The beads were resuspended in 50 mM NH_4_HCO_3_ supplemented with trypsin (Mass Spectrometry Grade, Promega; Madison, US) at an enzyme-to-protein ratio of 1:25 (w/w) and incubated overnight at 37°C. After overnight digestion, acetonitrile was added to the samples to reach a final concentration of 95% (v/v) followed by incubation at RT for 20 min. To increase the yield, supernatants derived from this initial peptide-binding step were additionally subjected to the SP3 peptide purification procedure (Sielaff et al., 2017). Each sample was washed with acetonitrile. To recover bound peptides, paramagnetic beads from the original sample and corresponding supernatants were pooled in 2% (v/v) dimethyl sulfoxide (DMSO) in water and sonicated for 1 min. After 2 min of centrifugation at 14,000xg and 4°C, supernatants containing tryptic peptides were transferred into a glass vial for MS analysis and acidified with 0.1% (v/v) formic acid.

### Liquid chromatography-mass spectrometry (LC-MS) analysis

Tryptic peptides were separated using an Ultimate 3000 RSLCnano LC system (Thermo Fisher Scientific; Waltham, US) equipped with a PEPMAP100 C18 5 µm 0.3 x 5 mm trap (Thermo Fisher Scientific; Waltham, US) and an HSS-T3 C18 1.8 μm, 75 μm x 250 mm analytical reversed-phase column (Waters Corporation; Milford, US). Mobile phase A was water containing 0.1% (v/v) formic acid and 3% (v/v) DMSO. Peptides were separated by running a gradient of 2–35% mobile phase B (0.1% (v/v) formic acid, 3% (v/v) DMSO in ACN) over 40 min at a flow rate of 300 nl/min. Total analysis time was 60 min including wash and column re-equilibration steps. Column temperature was set to 55°C. Mass spectrometric analysis of eluting peptides was conducted on an Orbitrap Exploris 480 (Thermo Fisher Scientific; Waltham, US) instrument platform. Spray voltage was set to 1.8 kV, the funnel RF level to 40, and heated capillary temperature was at 275°C. Data were acquired in data-dependent acquisition (DDA) mode targeting the 10 most abundant peptides for fragmentation (Top10). Full MS resolution was set to 120,000 at *m/z* 200 and full MS automated gain control (AGC) target to 300% with a maximum injection time of 50 ms. Mass range was set to *m/z* 350–1,500. For MS2 scans, the collection of isolated peptide precursors was limited by an ion target of 1 × 10^5^ (AGC tar-get value of 100%) and maximum injection times of 25 ms. Fragment ion spectra were acquired at a resolution of 15,000 at *m/z* 200. Intensity threshold was kept at 1E4. Isolation window width of the quadrupole was set to 1.6 *m/z* and normalized collision energy was fixed at 30%. All data were acquired in profile mode using positive polarity. Each sample was analyzed in three technical replicates.

### Data analysis and label-free quantification

DDA raw data acquired with the Exploris 480 were processed with MaxQuant (version 2.0.1) (Cox & Mann 2008; Cox et al., 2014), using the standard settings and label-free quantification (LFQ) enabled for each parameter group, i.e. control and affinity-purified samples (LFQ min ratio count 2, stabilize large LFQ ratios disabled, match-between-runs). Data were searched against the forward and reverse sequences of the *P. falciparum* proteome (UniProtKB/TrEMBL, 5,445 entries, UP000001450, released April 2020) and a list of common contaminants. For peptide identification, trypsin was set as protease allowing two missed cleavages. Carbamidomethylation was set as fixed and oxidation of methionine as well as acetylation of protein N-termini as variable modifications. Only peptides with a minimum length of 7 amino acids were considered. Peptide and protein false discovery rates (FDR) were set to 1%. In addition, proteins had to be identified by at least two peptides. Statistical analysis of the data was conducted using Student’s t-test, which was corrected by the Benjamini-Hochberg (BH) method for multiple hypothesis testing (FDR of 0.01). In addition, proteins in the affinity-enriched samples had to be identified in all three biological replicates and show at least a two-fold enrichment compared to the controls. The datasets of protein hits were further edited by verification of the gene IDs and gene names via the PlasmoDB database (www.plasmodb.org; Aurrecoechea et al., 2009). PlasmoDB IDs were extracted from the fasta headers provided by mass spectrometry and verified manually.

### Data availability

The mass spectrometry proteomics data have been deposited to the ProteomeXchange Consortium (http://proteomecentral.proteomexchange.org) via the jPOSTrepo partner repository (Okuda et al., 2017; Vizcaíno et al., 2013) with the dataset identifiers PXD039679 (ProteomeXchange) and JPST002011 (jPOSTrepo).

## Acknowledgements

The authors acknowledge funding by the Deutsche Forschungsgemeinschaft (Grants PR905/19-1 to GP, GI 312/11-1 to TWG and TE599/9-1 to ST of the DFG priority programme SPP 2225). The authors thank Pietro Alano (Istituto Superiore di Sanità Rome) for providing antisera against G377 and MDV1 and Ron Dzikowski (The Hebrew University of Jerusalem) for providing vector pSLI-HA-*glmS.* The authors further gratefully acknowledge the microscopy support from the Advanced Light and Fluorescence Microscopy (ALFM) facility at the Centre for Structural Systems Biology (CSSB).

## Notes

### Competing Interest Statement

The authors have declared no competing interest.

