## Supplemental Figures for "Comparative proteomics of vesicles essential for the egress of *Plasmodium falciparum* gametocytes from red blood cells"

*Original Research Article, Molecular Microbiology, Special Issue: Mechanisms of host cell exit by intracellular pathogens*

### Comparative proteomics of vesicles essential for the egress of *Plasmodium falciparum* gametocytes from red blood cells

Juliane Sassmannshausen[1], Sandra Bennink[1], Ute Distler[2], Juliane Küchenhoff[1], Allen M. Minns[3], Scott E. Lindner[3], Paul-Christian Burda[4,5,6], Stefan Tenzer[2], Tim W. Gilberger[4,5,6], and Gabriele Pradel[1]

#### Supplemental Figures S1-S10

**Figure S1A**

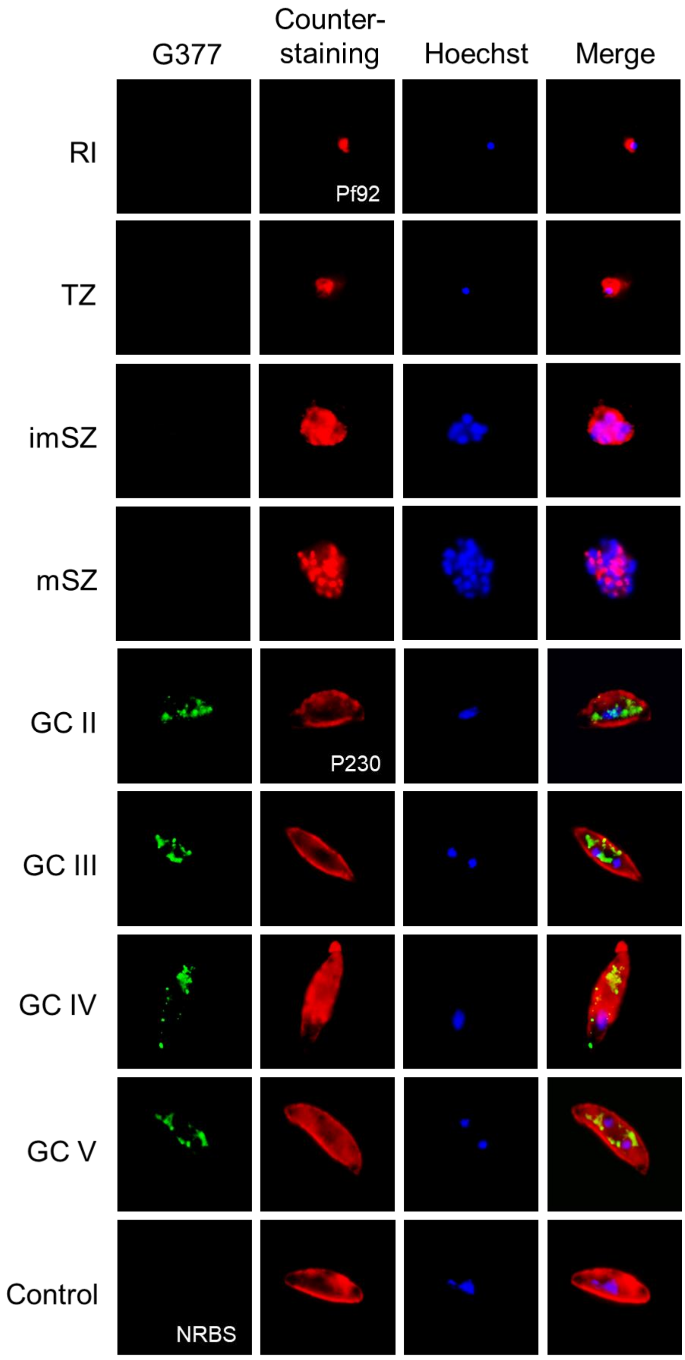

G377
Counter-staining
Hoechst
Merge
RI
Pf92
TZ
imSZ
mSZ
GC II
P230
GC III
GC IV
GC V
Control
NRBS

**Figure S1B**

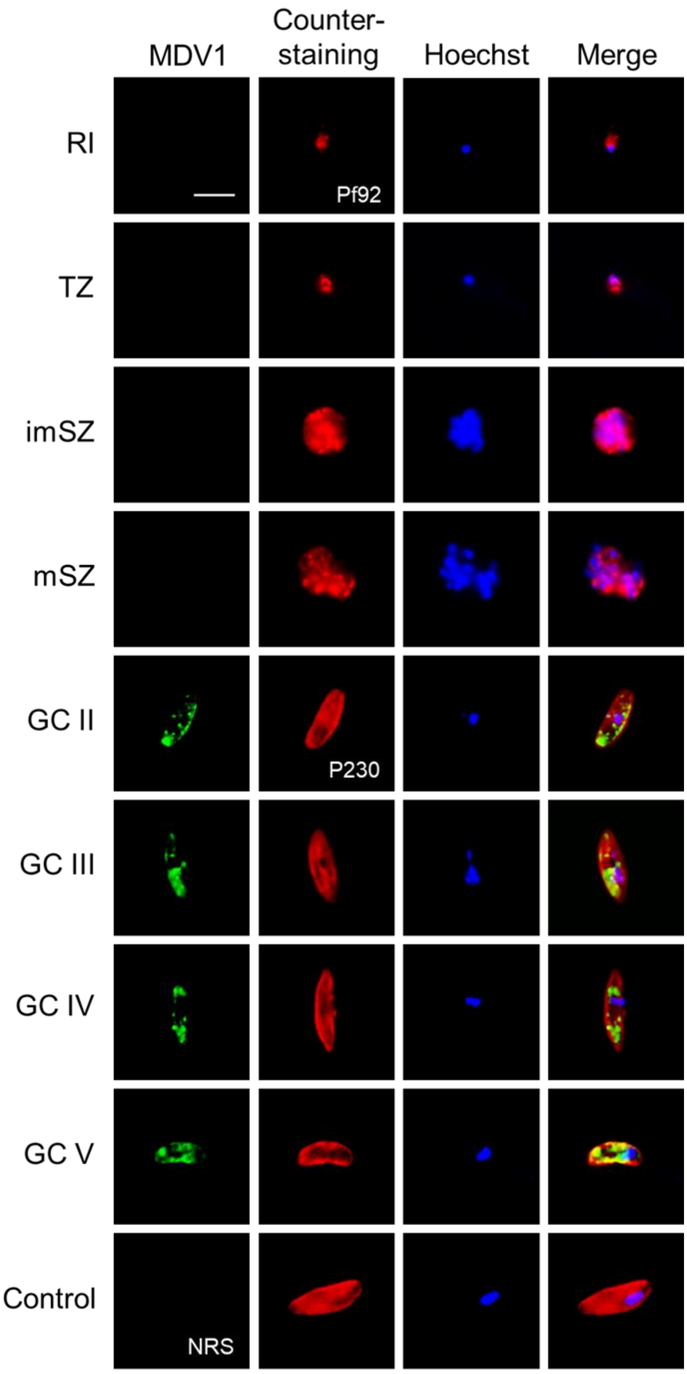
MDV1
Counter-
staining
Hoechst
Merge
RI
Pf92
TZ
imSZ
mSZ
GC II
P230
GC III
GC IV
GC V
Control
NRS

**Figure S1C**

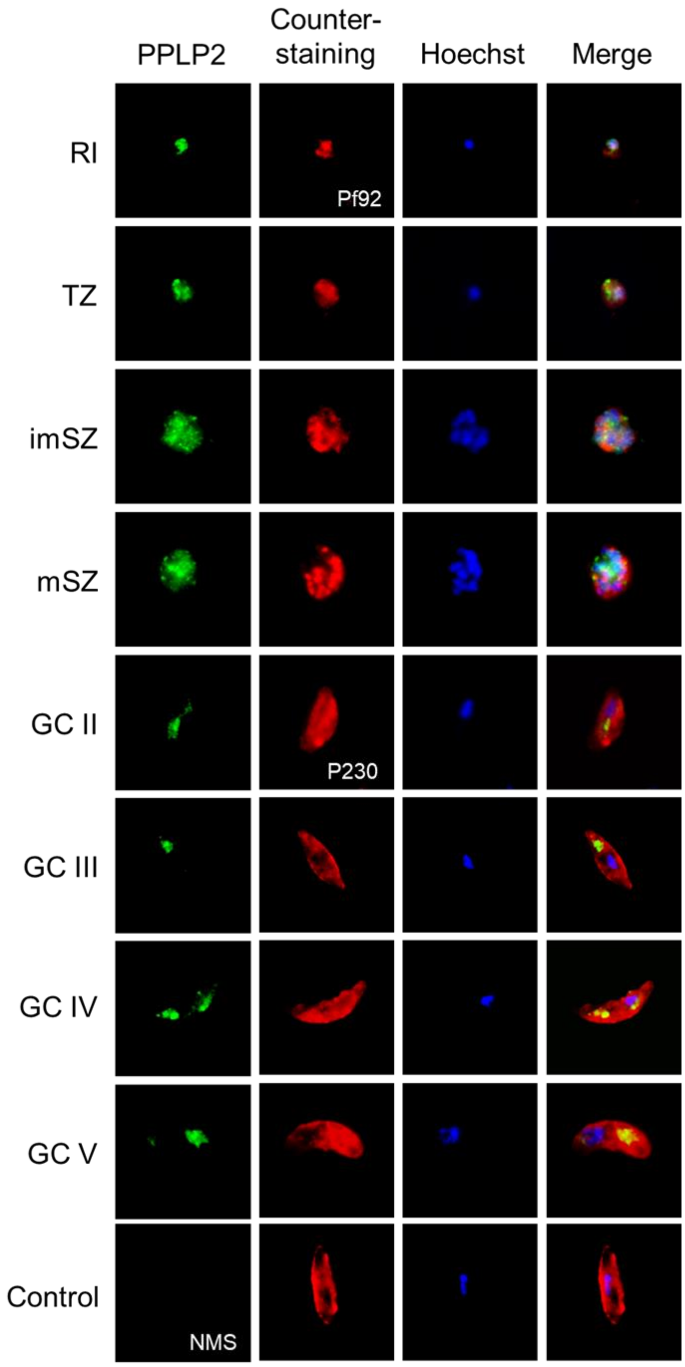


**Figure S1: Localization of G377, MDV1 and PPLP2 in the *P. falciparum* blood stages.** Asexual and sexual blood stages of WT NF54 were immunolabeled with rabbit anti-G377, rat anti-MDV1, and mouse anti-PPLP2 antisera, respectively (green) to highlight G377 **(A)**, MDV1 **(B)** and PPLP2 **(C)**. Asexual blood stages were counterstained by rabbit or mouse anti-

Pf92 antisera (red), gametocytes were counterstained by rabbit or mouse anti-P230 antisera (red); parasite nuclei were highlighted by Hoechst 33342 nuclear stain (blue). Antisera from non-immunized animals were used as negative control. Bar; 5 μm. RI, ring stage; TZ, trophozoite; imSZ, immature schizont; mSZ, mature schizont; GC II–V, gametocytes stages II to V; NRS, neutral rat serum; NRBS, neutral rabbit serum; NMS, neutral mouse serum. Results (A-C) are representative of three independent experiments.

**Figure S2**

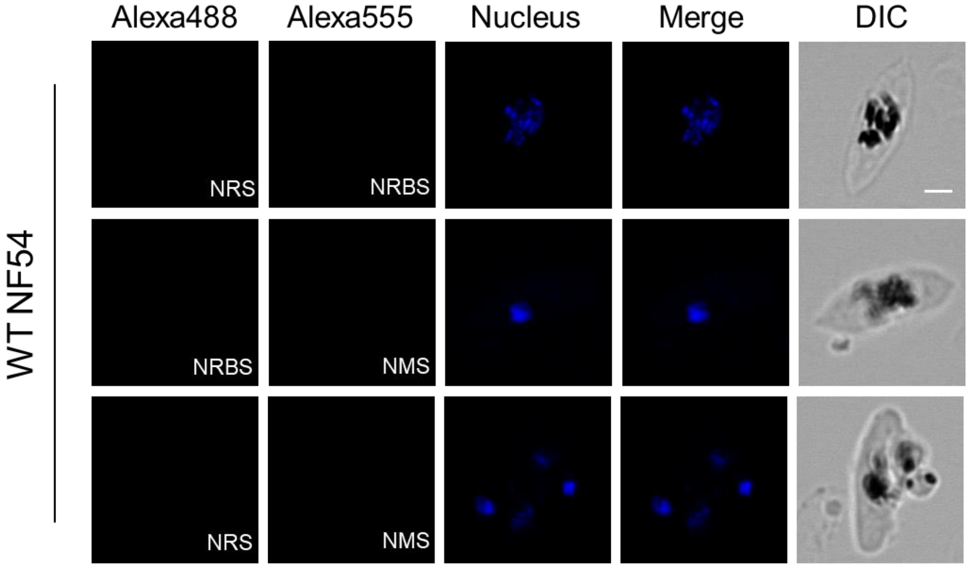


**Figure S2: IFA controls corresponding to Fig. 1B.** WT NF54 gametocytes were immunolabeled with antisera from non-immunized animals (green, Alexa488; red, Alexa555). Parasite nuclei were highlighted by Hoechst 33342 nuclear stain (blue). NRS, neutral rat serum; NRBS, neutral rabbit serum; NMS, neutral mouse serum; DIC, differential interference contrast. Bar; 2 µm. Results are representative of three independent experiments.

**Figure S3**

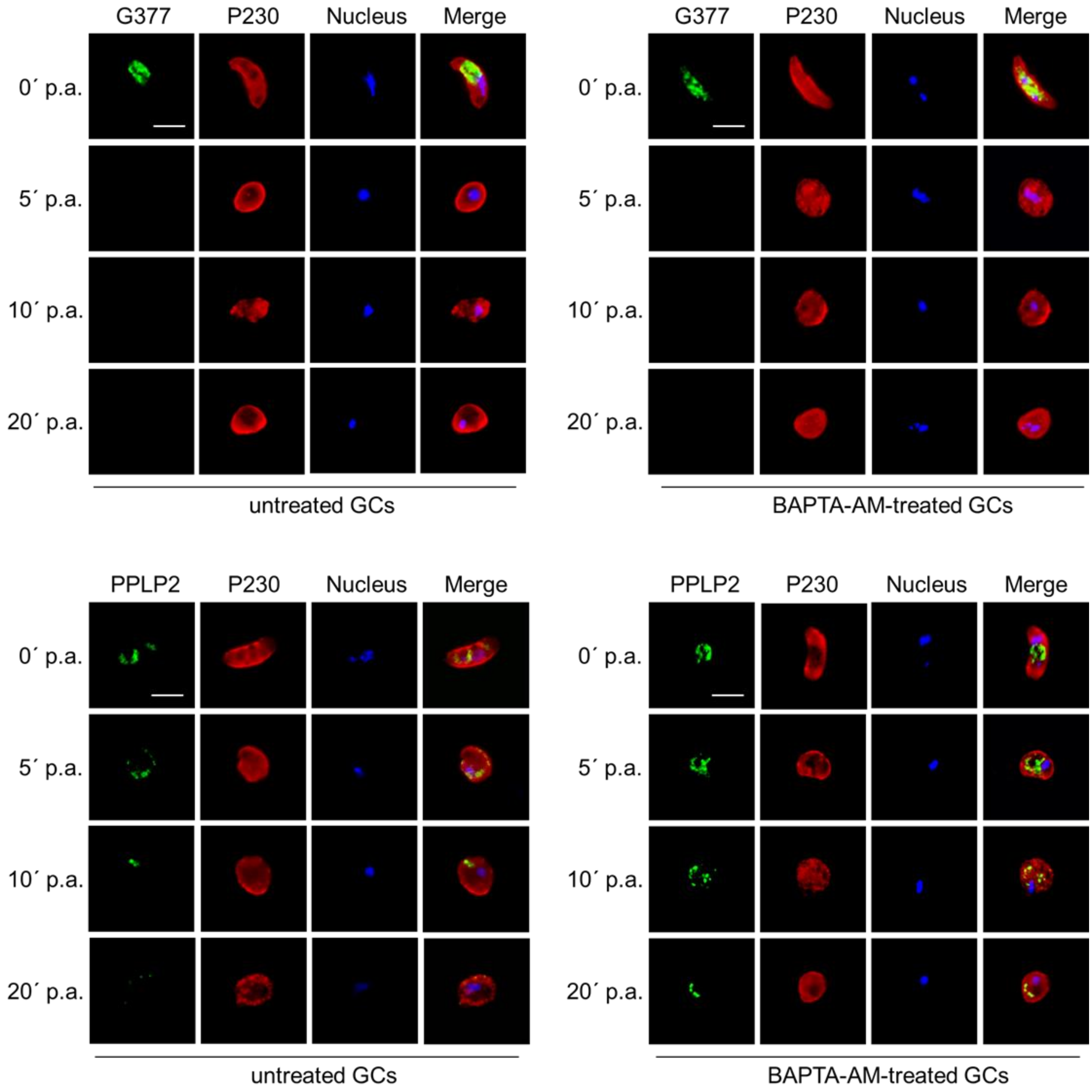


**Figure S3: Calcium-dependency of OB and g-exoneme discharge.** Mature WT NF54 gametocytes (GC) were treated with 25 µM BAPTA-AM for 15 min prior to activation. Untreated GC served as control. GCs were collected at 0-20 min post-activation (p.a.) and immunolabeled using rabbit anti-G377 or mouse anti-PPLP2 antisera (green). Gametocytes were counterstained with mouse or rabbit anti-P230 antisera (red). Parasite nuclei were highlighted by Hoechst 33342 nuclear stain (blue). Bar; 5 µm. Results are representative of three independent experiments.

**Figure S4**

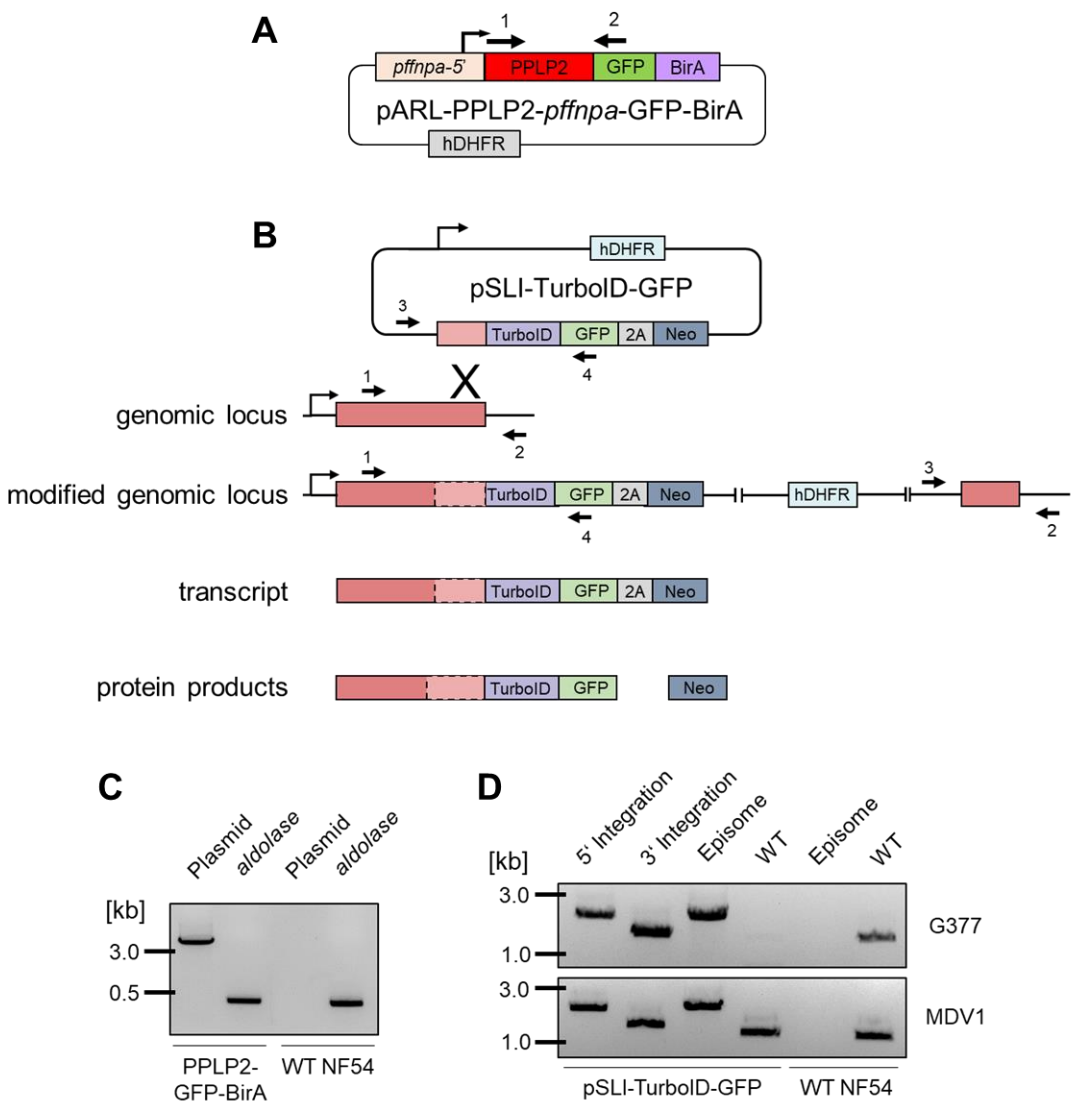


**Figure S4: Generation of the parasite lines to be used in BioID. (A)** Schematic depicting the vector pARL-PPLP2-*pffnpa*-BirA. **(B)** Schematic depicting the single-crossover homologous recombination strategy for the generation of the pSLI-TurboID-GFP-based transgenic lines. The coding region of the gene of interest was fused at the 3'-end to a sequence coding for an advanced *E. coli* biotin ligase and a GFP-encoding sequence followed by the 2A-skip peptide sequence and the Neo sequence. The numbered arrows indicate the positions of primers used to confirm vector integration. HA, hemagglutinin; hDHFR, human dihydrofolate reductase-encoding gene conferring resistance to WR99210; Neo, gene conferring resistance to neomycin. **(C)** Detection of episomal uptake PPLP2-GFP-BirA line. Diagnostic PCR was employed to detect the presence of vector pARL-PPLP2-*pffnpa*-GFP-BirA (primers 1 and 2) in the PPLP2-GFP-BirA line, but not the WT NF54. Primers specific for the gene encoding plasmodial aldolase were used as control. **(D)** Confirmation of vector integration into the gene locus coding for G377 (top) and MDV1 (bottom). Diagnostic PCR demonstrates successful 5′ (primers 1 and 4) and 3′ (primers 3 and 2) integration. As a control, WT NF54 gDNA was used, demonstrating the original gene locus (primers 1 and 2). Episomal DNA was further detected (primers 3 and 4). Primer sequences and band sizes are provided in Tables S4 and S5.

**Figure S5**

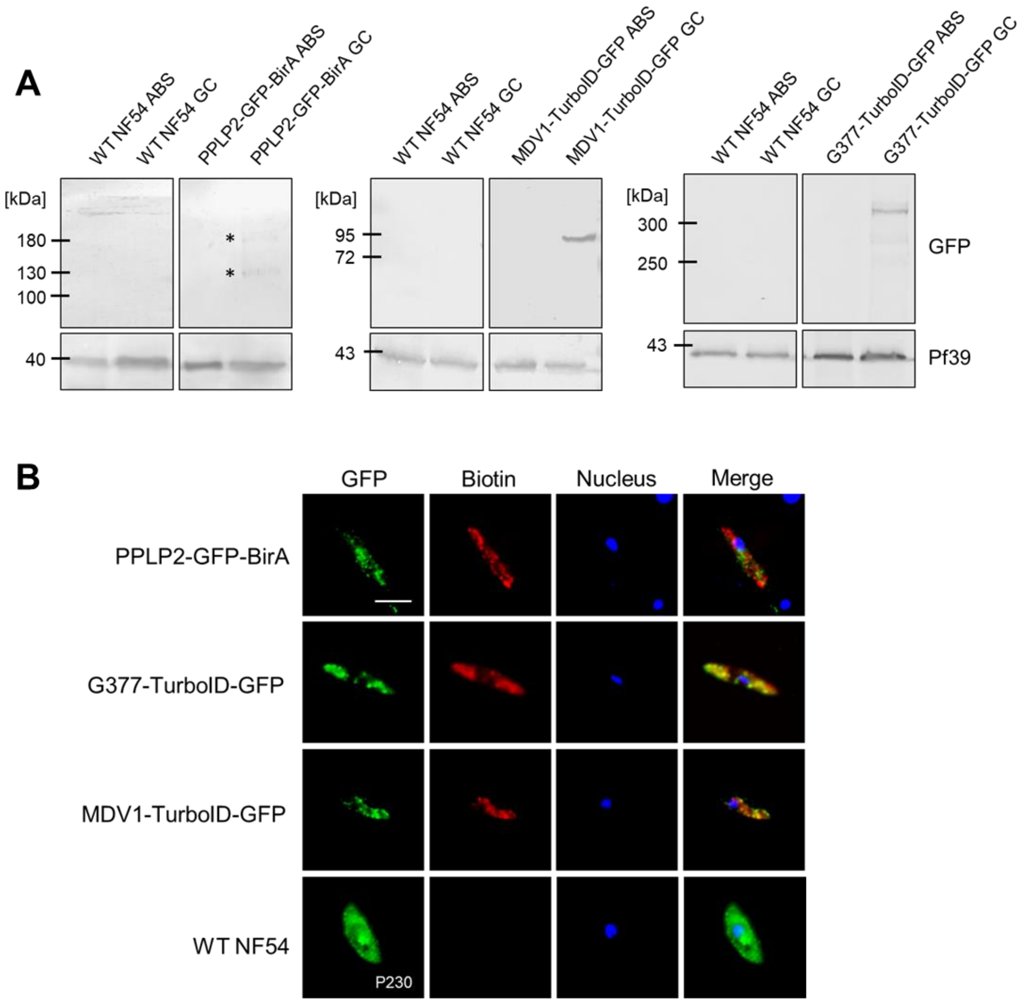


**Figure S5: Verification of the parasite lines to be used in BioID. (A)** Expression of the bait proteins in the BioID parasite lines. Lysates of asexual blood stages (ABS) and gametocytes (GC) of the PPLP2-GFP-BirA line and the G377-TurboID-GFP and MDV1-TurboID-GFP lines were immunoblotted with mouse anti-GFP antibody to detect the respective fusion protein (PPLP2-GFP-BirA, 187 kDa; MDV1-TurboID-GFP, 91 kDa; G377-TurboID-GFP, 439 kDa). ABS and GC lysate of WT NF54 served as negative controls, while immunoblotting with rabbit antibody against the ER protein Pf39 served as loading control (39 kDa). Asteriks (*) highlight the detected bands in the PPLP2-GFP-BirA GC lysate. **(B)** Localization of biotinylated proteins in the BioID parasite lines. Gametocytes of the PPLP2-GFP-BirA line and the G377-TurboID-GFP and MDV1-TurboID-GFP lines were treated with biotin (+) for 20 h (PPLP2-GFP-BirA) and 15 min (G377-, MDV-TurboID-GFP). Treated NF54 WT gametocytes served as controls.

Gametocytes of the transfectant lines were immunolabeled with mouse anti-GFP antibody to detect the respective fusion protein, WT NF54 gametocytes were labelled with rabbit anti-P230 sera (green). Biotinylated proteins were highlighted by fluorophore-conjugated streptavidin (red); parasite nuclei were highlighted by Hoechst 33342 nuclear stain (blue). Bar; 5 µm. Results (A, B) are representative of three independent experiments.

**Figure S6**

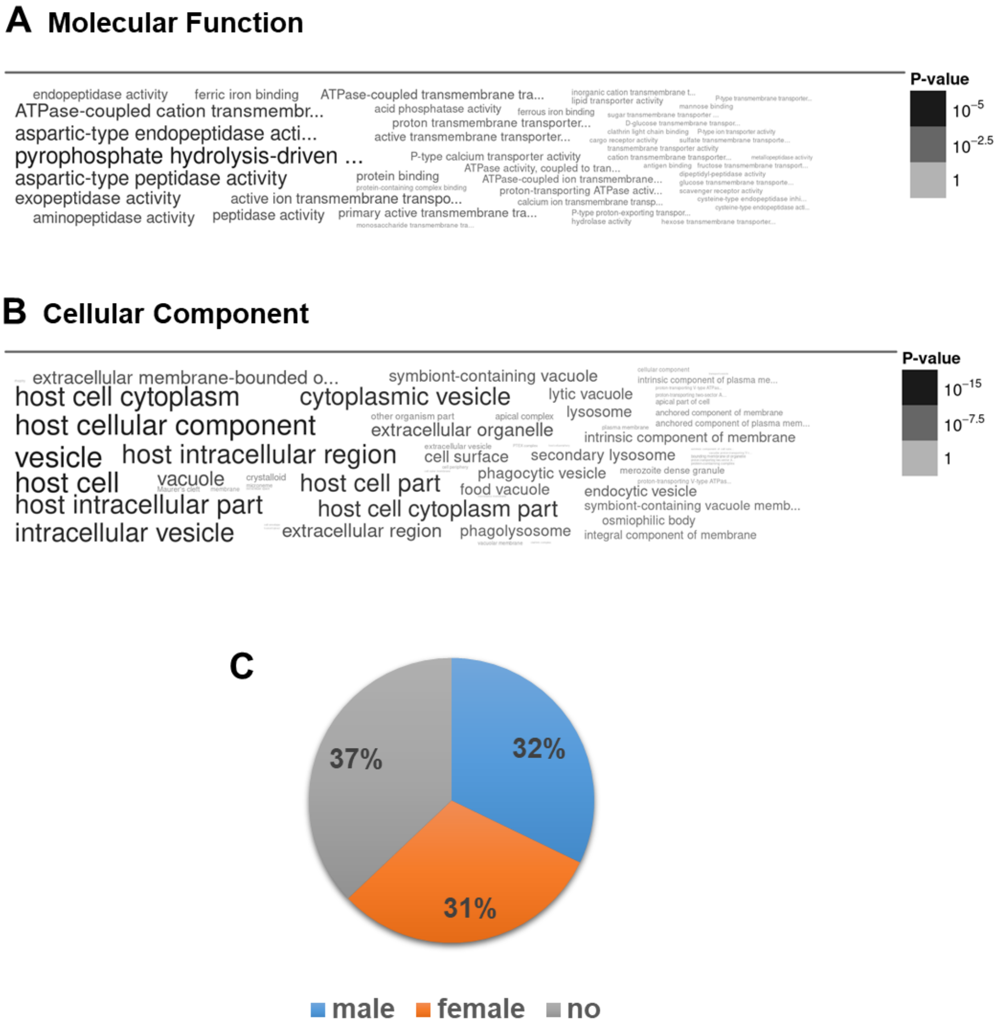


**Figure S6: GO analyses and sex specificity of egress vesicle proteins.** Word cloud visualizations of gene ontology enrichment (GO) analyses of the 143 egress vesicle proteins as identified by BioID analysis with respect to **(A)** molecular function and **(B)** cellular component. **(C)** Pie chart depicting the egress vesicle proteins (percentage of total numbers) grouped by sex (Lasonder et al., 2016; see PlasmoDB database).

**Figure S7**

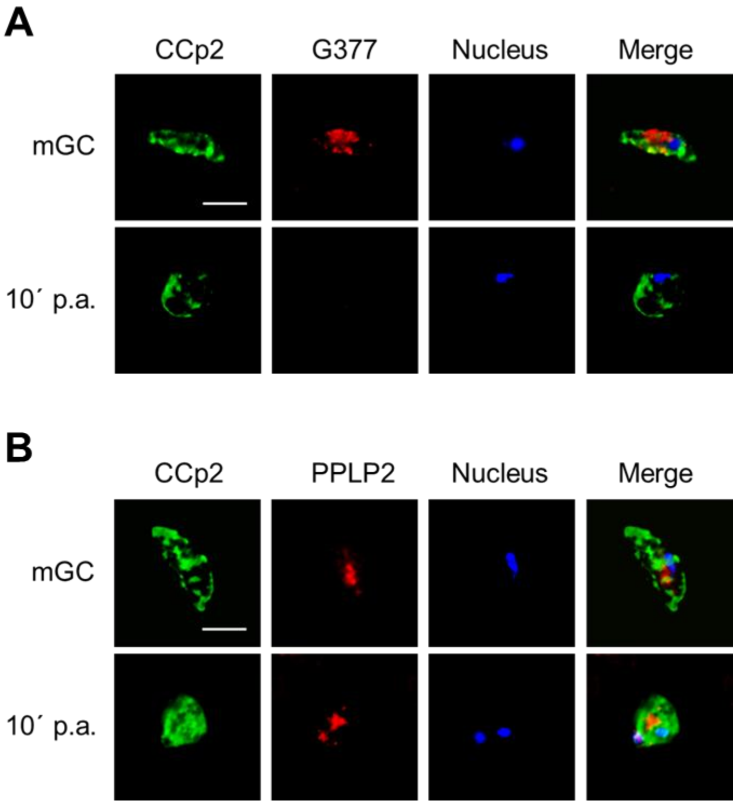


**Figure S7: Localization of the LCCL domain protein CCp2 in gametocytes.** WT NF54 gametocytes were collected at 0 and 10 min post-activation (p.a.) and immunolabeled with mouse or rabbit anti-CCp2 antisera (green). OBs **(A)** and g-exonemes **(B)** were highlighted by rabbit anti-G377 and mouse anti-PPLP2 antisera (red); parasite nuclei were highlighted by Hoechst 33342 nuclear stain (blue). Bar; 5 µm. Results (A, B) are representative of three independent experiments.

**Figure S8**

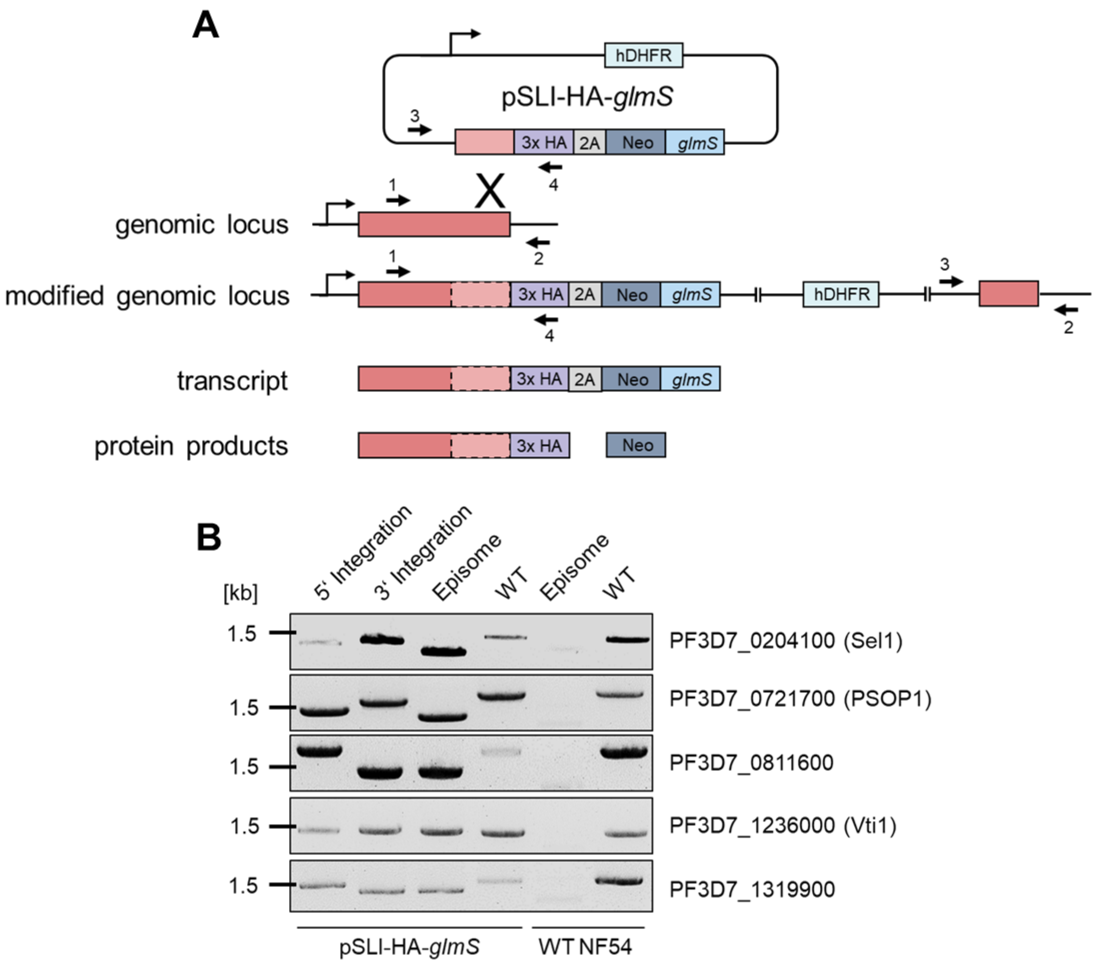


**Figure S8: Generation of pSLI-HA-*glmS*-based parasite lines. (A)** Schematic depicting the single-crossover homologous recombination strategy for the generation of the pSLI-HA-*glmS*-based transfectant lines. The coding region of the gene of interest was fused at the 3'-end to a HA-encoding sequence followed by the 2A-skip peptide sequence and the Neo and *glmS*-ribozyme sequences. The numbered arrows indicate the positions of primers used to confirm vector integration. *glmS*, glucosamine-6-phosphate-activated ribozyme; HA, hemagglutinin; hDHFR, human dihydrofolate reductase-encoding gene conferring resistance to WR99210; Neo, gene conferring resistance to neomycin. **(B)** Confirmation of vector integration into the gene locus coding for the respective egress vesicle protein. Diagnostic PCR demonstrates successful 5′ (primers 1 and 4) and 3′ (primers 3 and 2) integration. As a control, WT NF54 gDNA was used, demonstrating the original gene locus (primers 1 and 2). Episomal DNA was further detected (primers 3 and 4). Primer sequences and band sizes are provided in Tables S4 and S5.

**Figure S9**

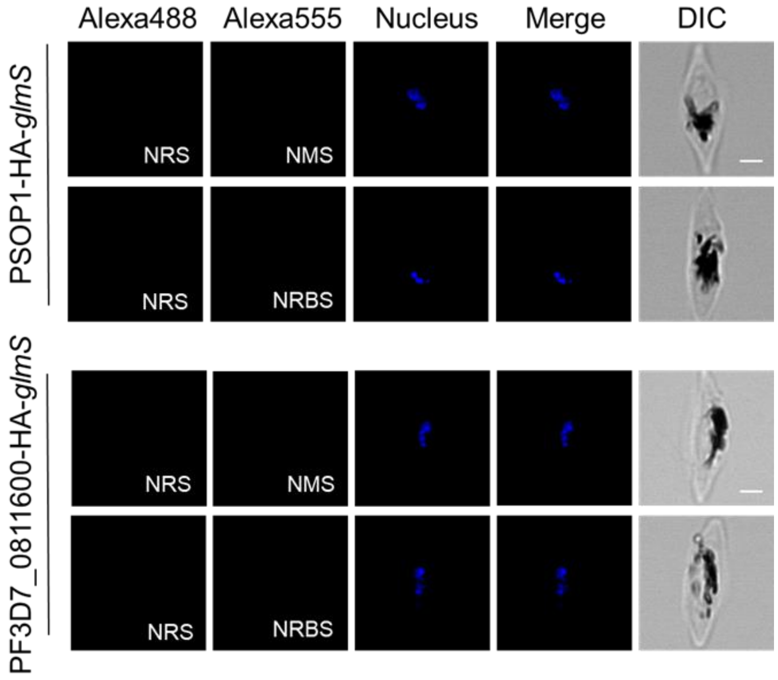


**Figure S9: IFA controls corresponding to Fig. 6A.** Gametocytes of lines PSOP1-HA-*glmS* and PF3D7_0811600-HA-*glmS* were immunolabeled with antisera from non-immunized animals (green, Alexa488; red, Alexa555). Parasite nuclei were highlighted by Hoechst 33342 nuclear stain (blue). NRS, neutral rat serum; NRBS, neutral rabbit serum; NMS, neutral mouse serum; DIC, differential interference contrast. Bar; 2 µm. Results are representative of three independent experiments.

**Figure S10**

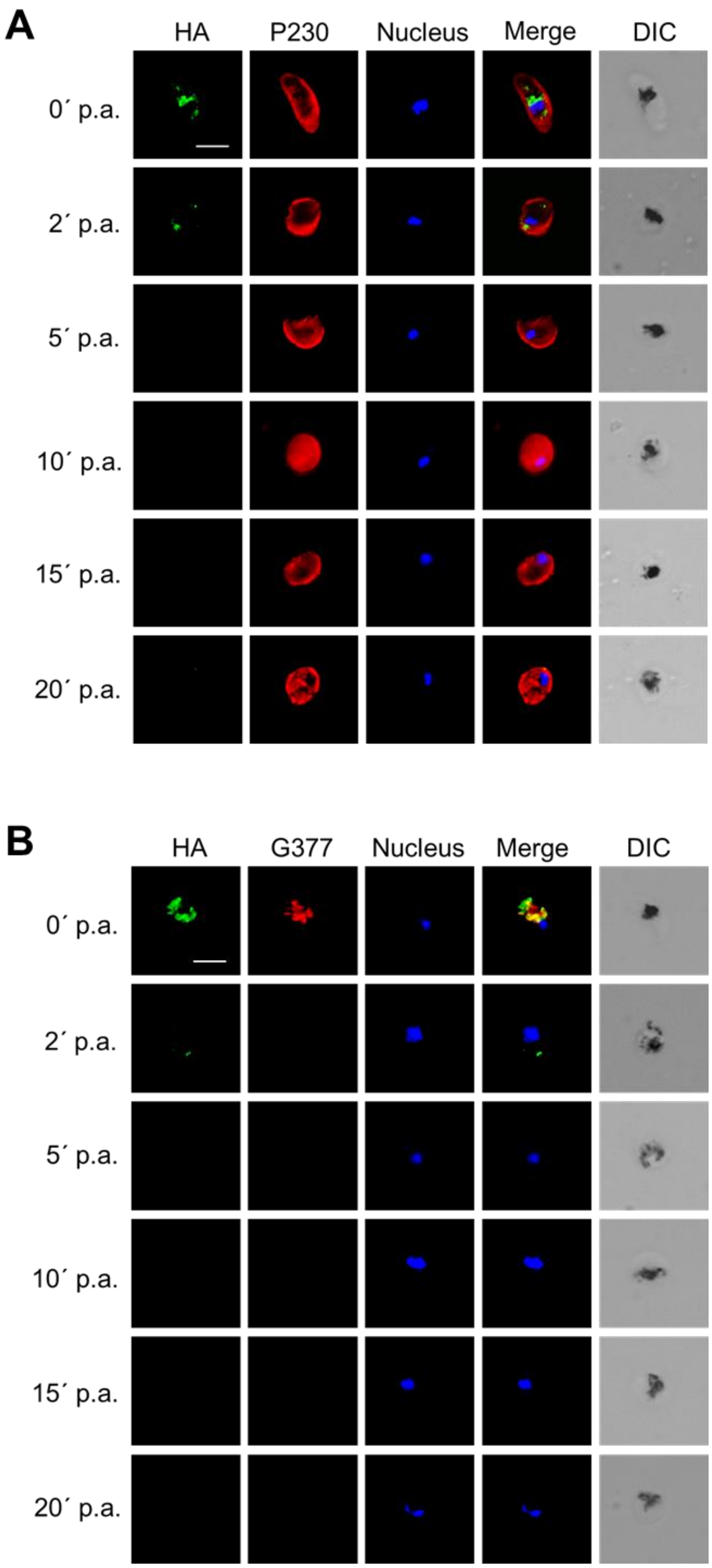
A
HA
P230
Nucleus
Merge
DIC
0´ p.a.
2´ p.a.
5´ p.a.
10´ p.a.
15´ p.a.
20´ p.a.
B
HA
G377
Nucleus
Merge
DIC
0´ p.a.
2´ p.a.
5´ p.a.
10´ p.a.
15´ p.a.
20´ p.a.

**Figure S10: PSOP1 discharge following gametocyte activation.** Gametocytes of line PSOP1-HA-*glmS* were collected at 0-20 min post-activation (p.a.) and immunolabeled with rat anti-HA antibody to highlight PSOP1 (green). **(A)** Gametocytes were counterstained by rabbit anti-P230 antisera (red), or **(B)** OBs were highlighted by rabbit anti-G377 antisera (red); parasite nuclei were highlighted by Hoechst 33342 nuclear stain (blue). Bar; 5 µm. DIC, differential interference contrast. Results (A, B) are representative of three independent experiments.
