## Supplemental Table S4 + S5 for "Comparative proteomics of vesicles essential for the egress of *Plasmodium falciparum* gametocytes from red blood cells"

*Original Research Article, Molecular Microbiology, Special Issue: Mechanisms of host cell exit by intracellular pathogens*

**Comparative proteomics of vesicles essential for the egress of *Plasmodium falciparum* gametocytes from red blood cells**

Juliane Sassmannshausen[1], Sandra Bennink[1], Ute Distler[2], Juliane Küchenhoff[1], Allen M. Minns[3], Scott E. Lindner[3], Paul-Christian Burda[4,5,6], Stefan Tenzer[2], Tim W. Gilberger[4,5,6], and Gabriele Pradel[1]

### Supplemental Tables S4 & S5

**Table S4. List of primers.**

| Function | Name | Sequence 5‘-3‘ |
|---|---|---|
| **Oligonucleotides for the generation and verification of the PPLP2-GFP-BirA line** | | |
| Cloning Primers | PPLP2 BirA KpnI FP | aaaaaGGTACCATGAATATACTCCTCAAATATC |
| | PPLP2 BirA AvrII RP | aaaaaCCTAGGTTCCACCAAATTGTTTGCCCC |
| Diagnostic PCR primers | PPLP2 BirA KpnI FP | aaaaaGGTACCATGAATATACTCCTCAAATATC |
| | pGREP RP | GCATTGAAGACCATACGCGAAAGTAGTGAC |
| | Aldolase FP | TAGATGGATTAGCAGAAAGATGC |
| | Aldolase RP | TCAAAATCACGCACATCCTG |
| **Oligonucleotides for the generation and verification of the G377-TurboID-GFP line** | | |
| Cloning primers | G377 TurboID NotI FP | aaaaagcggccgcCTCAACGGAAATTATTTGAGT |
| | G377 TurboID SpeI RP | aaaaaACTAGTTAATTCGTAAAGGTCTAGACC |
| Integration PCR primers | 5´ Int G377 TurboID (1) | TGTTCATGTCTATTC |
| | 3´ Int G377 TurboID (2) | CTTTTCACCAACACTTAA |
| | pARL-HA-*glmS* FP (3) | GCTTTACACTTTATGCTTCCGGCTCG |
| | pGREP RP (4) | GCATTGAAGACCATACGCGAAAGTAGTGAC |
| **Oligonucleotides for the generation and verification of the MDV1-TurboID-GFP line** | | |
| Cloning Primers | MDV1 TurboID NotI FP | aaaaaGCGGCCGCCAGAGTCATGGGAATTTA |
| | MDV1 TurboID SpeI RP | aaaaaACTAGTATCACTATCACTGTGTGT |
| Integration PCR Primers | 5´ Int MDV1 TurboID (1) | CTGAATGGAGTAATAGAG |
| | 3´ Int MDV1 TurboID (2) | ATACCGTCTATATGAGTC |
| | pARL-HA-*glmS* FP (3) | GCTTTACACTTTATGCTTCCGGCTCG |
| | pGREP RP (4) | GCATTGAAGACCATACGCGAAAGTAGTGAC |
| **Oligonucleotides for the generation and verification of the PF3D7_0204100/Sel1-HA-*glmS* line** | | |
| Cloning Primers | 0204100 pSLI *glmS* NotI FP | atgcatGCGGCCGCTAAACATTTACGGATTAGTGAG |
| | 0204100 pSLI *glmS* XmaI RP | atgcatCCCGGGTATAATTTTAAAAAAAGATAATATCAG |
| | 5´ Int 0204100 pSLI-*glmS* (1) | ATGAGAATGTACAACGTG |

| | | |
|---|---|---|
| Integration PCR Primers | 3´ Int 0204100 pSLI-*glmS* (2) | CCAACCCTAATTAACAC |
| | pARL-HA-*glmS* FP (3) | GCTTTACACTTTATGCTTCCGGCTCG |
| | pSLI-HA-*glmS* RP (4) | TGTCTGTTGTGCCCAGTCAT |
| **Oligonucleotides for the generation and verification of the PF3D7_0721700/PSOP1-HA-*glmS* line** | | |
| Cloning Primers | 0721700 pSLI *glmS* NotI FP | atgcatGCGGCCGCTAAAGCTCCATATCAACTACC |
| | 0721700 pSLI *glmS* XmaI RP | atgcatCCCGGGTGGACTCCTTACAAATGAGG |
| Integration PCR Primers | 5´ Int 0721700 pSLI-*glmS* (1) | ACGTTGATGCCAATGGAG |
| | 3´ Int 0721700 pSLI-*glmS* (2) | GACCGATATAGTGCTCTCC |
| | pARL-HA-*glmS* FP (3) | GCTTTACACTTTATGCTTCCGGCTCG |
| | pSLI-HA-*glmS* RP (4) | TGTCTGTTGTGCCCAGTCAT |
| **Oligonucleotides for the generation and verification of the PF3D7_0811600-HA-*glmS* line** | | |
| Cloning Primers | 0811600 pSLI *glmS* NotI FP | atgcatGCGGCCGCTAAATGATGATATTAGTCTTGG |
| | 0811600 pSLI *glmS* XmaI RP | atgcatCCCGGGTTCGTCTACCTTAAATAAATAAAAGAAGACG |
| Integration PCR Primers | 5´ Int 0811600 pSLI-*glmS* (1) | AGTGCAGATGTATATAAGAG |
| | 3´ Int 0811600 pSLI-*glmS* (2) | TATAGGATACAAATAATTACGTG |
| | pARL-HA-*glmS* FP (3) | GCTTTACACTTTATGCTTCCGGCTCG |
| | pSLI-HA-*glmS* RP (4) | TGTCTGTTGTGCCCAGTCAT |
| **Oligonucleotides for the generation and verification of the PF3D7_1236000/Vti1-HA-*glmS* line** | | |
| Cloning Primers | 1236000 pSLI *glmS* NotI FP | ATCGATGCGGCCGCTAACTGGAGGTCGAAGC |
| | 1236000 pSLI *glmS* XmaI RP | ATCGATCCCGGGTGAGGAGGTGT |
| Integration PCR Primers | 5´ Int 1236000 pSLI-*glmS* (1) | CACGGAGAAGATAATTAAGG |
| | 3´ Int 1236000 pSLI-*glmS* (2) | ACACATGTCGAAATGTATGC |
| | pARL-HA-*glmS* FP (3) | GCTTTACACTTTATGCTTCCGGCTCG |
| | pSLI-HA-*glmS* RP (4) | TGTCTGTTGTGCCCAGTCAT |
| **Oligonucleotides for the generation and verification of the PF3D7_1319900-HA-*glmS* line** | | |
| Cloning Primers | 1319900 pSLI *glmS* NotI FP | atgcatGCGGCCGCTAAACGTTAATGAATGTAGC |

| | | |
|---|---|---|
| | 1319900 pSLI *glmS* XmaI RP | atgcatCCCGGGTTCGTTTATGTAAAATACATTAAGG |
| Integration PCR Primers | 5´ Int 1319900 pSLI-*glmS* (1) | ACATTCAAATTCACATTCCG |
| | 3´ Int 1319900 pSLI-*glmS* (2) | ATATGTGCATTGAGTGAGC |
| | pARL-HA-*glmS* FP (3) | GCTTTACACTTTATGCTTCCGGCTCG |
| | pSLI-HA-*glmS* RP (4) | TGTCTGTTGTGCCCAGTCAT |

Numbers in brackets indicate primer numbers (compare Figs. S4 and S8).

FP, forward primer; RP, reverse primer

**Table S5. List of integration PCR primer combinations and the expected fragment sizes.**

| Fragment | Primer combinations | Size [bp] |
|---|---|---|
| **PPLP2 -GFP-BirA** | | |
| Episome | PPLP2 BirA KpnI FP + pGREP RP | 3414 |
| Aldolase | Aldolase FP + Aldolase RP | 378 |
| **G377-TurboID-GFP** | | |
| 5´ Integration | 5´Int G377 TurboID (1) + pGREP RP (4) | 2214 |
| 3´ Integration | pARL-HA-*glmS* FP (3) + 3´Int G377 TurboID (2) | 1196 |
| Episome | 5´Int G377 TurboID (1) + 3´Int G377 TurboID (2) | 2156 |
| Wildtype | pARL-HA-*glmS* FP (3) + pGREP RP (4) | 1254 |
| **MDV1-TurboID-GFP** | | |
| 5´ Integration | 5´Int MDV1 TurboID (1) + pGREP RP (4) | 1682 |
| 3´ Integration | pARL-HA-*glmS* FP (3) + 3´Int MDV1 TurboID (2) | 850 |
| Episome | 5´Int MDV1 TurboID (1) + 3´Int MDV1 TurboID (2) | 1757 |
| Wildtype | pARL-HA-*glmS* FP (3) + pGREP RP (4) | 775 |
| **PF3D7_0204100-HA-*glmS*** | | |
| 5´ Integration | 5´Int 0204100 pSLI-*glmS* (1) + pSLI-HA-*glmS* RP (4) | 1696 |
| 3´ Integration | pARL-HA-*glmS* FP (3) + 3´Int 0204100 pSLI-*glmS* (2) | 1726 |
| Episome | pARL-HA-*glmS* FP (3) + pSLI-HA-*glmS* RP (4) | 1277 |
| Wildtype | 5´Int 0204100 pSLI-*glmS* (1) + 3´Int 0204100 pSLI-*glmS* (2) | 2141 |
| **PF3D7_0721700-HA-*glmS*** | | |
| 5´ Integration | 5´Int 0721700 pSLI-*glmS* (1) + pSLI-HA-*glmS* RP (4) | 1447 |
| 3´ Integration | pARL-HA-*glmS* FP (3) + 3´Int 0721700 pSLI-*glmS* (2) | 1646 |
| Episome | pARL-HA-*glmS* FP (3) + pSLI-HA-*glmS* RP (4) | 1085 |
| Wildtype | 5´Int 0721700 pSLI-*glmS* (1) + 3´Int 0721700 pSLI-*glmS* (2) | 2008 |
| **PF3D7_0811600-HA-*glmS*** | | |
| 5´ Integration | 5´Int 0811600 pSLI-*glmS* (1) + pSLI-HA-*glmS* RP (4) | 1865 |
| 3´ Integration | pARL-HA-*glmS* FP (3) + 3´Int 0811600 pSLI-*glmS* (2) | 1267 |
| Episome | pARL-HA-*glmS* FP (3) + pSLI-HA-*glmS* RP (4) | 1281 |
| Wildtype | 5´Int 0811600 pSLI-*glmS* (1) + 3´Int 0811600 pSLI-*glmS* (2) | 1856 |
| **PF3D7_1236000-HA-*glmS*** | | |
| 5´ Integration | 5´Int 1236000 pSLI-*glmS* (1) + pSLI-HA-*glmS* RP (4) | 1404 |
| 3´ Integration | pARL-HA-*glmS* FP (3) + 3´Int 1236000 pSLI-*glmS* (2) | 1416 |
| Episome | pARL-HA-*glmS* FP (3) + pSLI-HA-*glmS* RP (4) | 1441 |
| Wildtype | 5´Int 1236000 pSLI-*glmS* (1) + 3´Int 1236000 pSLI-*glmS* (2) | 1394 |
| **PF3D7_1319900-HA-*glmS*** | | |

| | | |
|---|---|---|
| 5´ Integration | 5´Int 1319900 pSLI-*glmS* (1) + pSLI-HA-*glmS* RP (4) | 1419 |
| 3´ Integration | pARL-HA-*glmS* FP (3) + 3´Int 1319900 pSLI-*glmS* (2) | 1220 |
| Episome | pARL-HA-*glmS* FP (3) + pSLI-HA-*glmS* RP (4) | 1203 |
| Wildtype | 5´Int 1319900 pSLI-*glmS* (1) + 3´Int 1319900 pSLI-*glmS* (2) | 1436 |

Numbers in brackets indicate primer numbers (compare Figs. S4 and S8).
